# Decoupling AMPK from fatty acid synthesis allows maintenance of fitness late in life

**DOI:** 10.1101/2025.03.27.645766

**Authors:** Hanane Hadj-Moussa, Megan Ulusan, Dorottya Horkai, Mohammed Kamran Afzal Mirza, Jonathan Houseley

## Abstract

Although lifespan has long been the focus of ageing research, preventing functional decline late in life is a more pressing societal need. Here, we investigate the basis of senescence and declining fitness during replicative ageing in budding yeast, and describe a metabolic perturbation that preserves late-life fitness even on an unrestricted glucose diet. We show that senescence can be prevented by constitutive activation of AMPK, though only for approximately half the ageing population, and use genetic and functional assays to link this heterogeneous response with differences in cytosolic acetyl coenzyme A (Acetyl-CoA) metabolism. In one class of ageing cell, AMPK activity maintains fitness late-in-life through pathways that transport cytosolic Acetyl-CoA into mitochondria, but AMPK also inhibits fatty acid synthesis which leads to lipid starvation in the other class of ageing cell. Therefore, AMPK activity has both positive and negative effects, but we show that constitutive AMPK activity uncoupled from fatty acid synthesis inhibition (the A2A mutant) suppresses senescence and maintains fitness in both classes of ageing cell. Our findings support a model in which lipid starvation and excess acetyl coenzyme A availability are major drivers of senescence in replicatively aged wild-type yeast. This work shows that ageing is not intrinsically associated with declining fitness, at least in yeast, and that re-engineering highly conserved metabolic pathways allows fitness to be preserved very late in life.

**Graphical abstract:** 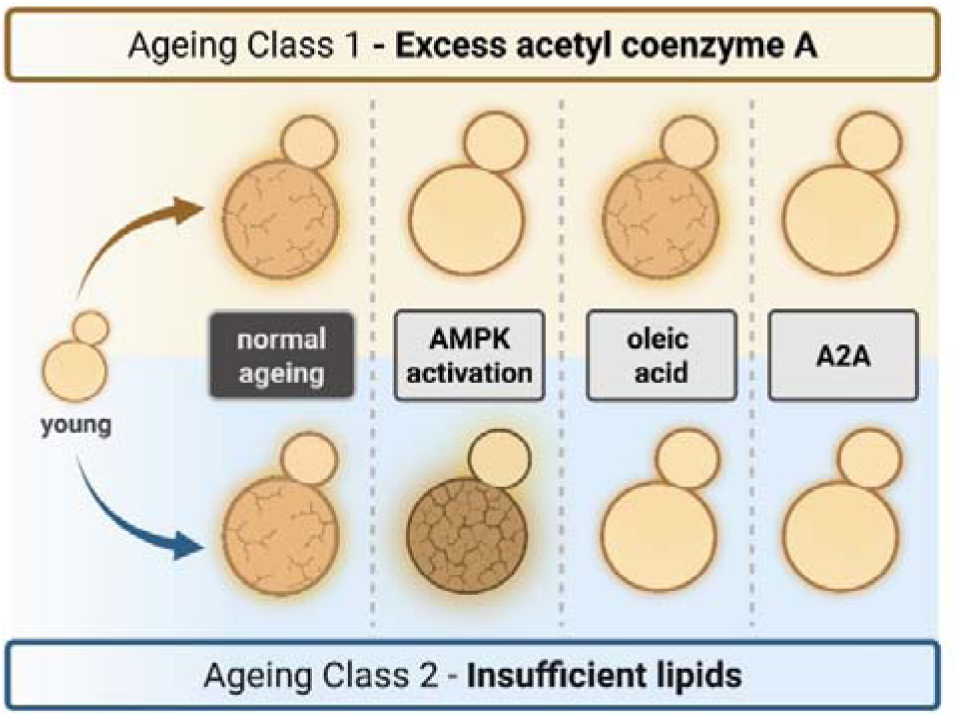

## Introduction

Traditional conceptions of ageing implicate a progressive accumulation of damage leading to systemic degradation in performance until death, with evolutionary pressures acting to maximise early life fitness and fecundity at the expense of ageing health, forming an antagonistic pleiotropy (1–3). The healthspan problem – the societal need to increase the fraction of human life spent in good health – is intractable under such models, since poor health late in life is an inevitable intermediate stage in the damage accumulation leading to death. However, ageing is not uniform across populations and from yeast to humans some members of any population maintain excellent fitness late in life (4–6); increasing the population fraction of such low-senescence individuals is a primary aim of ageing research. Interventions such as caloric restriction and mTOR inhibition can improve late-life fitness (7–11), but caloric restriction is an unrealistic societal intervention and mTOR inhibition has side effects, so new interventions are required.

Replicative ageing of the budding yeast *Saccharomyces cerevisiae* is a widely used model of ageing. Budding yeast cells divide asymmetrically into a mother cell and a daughter cell, with the mother cell undergoing only a limited number of divisions before permanently exiting the cell cycle (12). Mother cells maintain a rapid and uniform cell cycle for most of their replicative lifespan but the cell cycle usually slows dramatically in the last few divisions indicating loss of fitness (12–14), an effect known as ‘senescence’ following the classical definition of the term ‘senescence’ to denote a period of declining fitness for the organism at the end of life. This should not be equated with cellular senescence, the permanent cell cycle arrest caused by irreparable damage in mammalian cells. A sharp transition termed the senescence entry point (SEP) marks the change from rapid to slow, heterogeneous cycling and coincides with the appearance of apparent pathological markers (14,15). Not all cells senesce, and populations are extremely heterogeneous in some experimental systems, with individual cells following either a short-lived trajectory marked by an early SEP and accumulated foci of a mitochondrial Tom70-GFP marker when present, or a long-lived trajectory with little evidence of senescence (6,16,17). We recently reported that ageing yeast on galactose instead of glucose prevented senescence without extending lifespan across the whole population, providing a massive improvement in late-life fitness without caloric restriction (18).

The relevant metabolic differences caused by ageing on galactose are unclear, but suppression of senescence required both respiration and AMPK (18), a highly conserved kinase that reconfigures energy utilisation in response to poor nutrient availability by increasing respiration, mobilising internal energy stores, inducing autophagy and moderating fatty acid biosynthesis (reviewed in (19,20)). Due to its pivotal role in the response to low energy, AMPK has long been a focus of efforts to understand caloric restriction (21,22) and is required for the benefits of caloric restriction in yeast, worms and flies (23–25). Overexpression of the AMPK catalytic subunit extends lifespan in *C. elegans* and *D. melanogaster*, and high doses of the indirect AMPK activator metformin have been reported to extend lifespan and healthspan of mice (26–30). However, the mechanism underlying these lifespan effects is unclear, and the generality of these effects is uncertain, as other studies of metformin have reported no lifespan improvement in flies, mice or humans (31,32).

Caloric restriction robustly extends lifespan of budding yeast (33–35), but AMPK activity has different effects on yeast lifespan depending on the ageing paradigm employed, shortening replicative lifespan but extending chronological lifespan (36,37). Yeast AMPK is a central regulator of energy metabolism and shares close mechanistic and functional parallels with its counterpart in higher eukaryotes. Snf1, the catalytic subunit of AMPK in yeast, is activated by phosphorylation at a structurally conserved site, Thr210, by kinases Sak1, Elm1 and Tos3, and is regulated in response to glucose availability through Snf1 dephosphorylation, pH and subunit binding (38–40).

Suppression of fatty acid synthesis through inhibitory phosphorylation of acetyl coenzyme A carboxylase (ACC) is a highly conserved function of AMPK; this has been described for yeast Acc1, *Drosophila* ACC and mammalian ACC1/ACC2 (41–44). Fatty acids are synthesised from cytosolic Acetyl-CoA, which ultimately derives from glycolysis either through respiration in mammals or through oxidation of fermentation intermediates to acetate in yeast. Regardless of the source, fatty acid synthesis above immediate requirements acts as a sink of energy and carbon that can be mobilised during future scarcity by AMPK through suppression of fatty acid synthesis and promotion of fatty acid catabolism.

Here, we show that senescence and fitness loss in replicatively ageing yeast can be almost completely avoided without extension of lifespan, by rewiring the conserved AMPK-fatty acid metabolic regulatory system.

## Results

### AMPK suppresses senescence through mitochondrial Acetyl-CoA import pathways

AMPK is active and necessary for growth on galactose but not on glucose (45,46) and is required for cells ageing on galactose to avoid senescence (18). We therefore asked whether activation of AMPK during ageing on glucose can recapitulate the benefits of ageing on galactose. For this, we compared log phase populations of *S. cerevisiae* (average age 0-1 generation) to purified cohorts aged on rich glucose media for 24 or 48 hours (average replicative ages ∼12 and ∼18 generations, at which points cells are ∼95% and ∼30% viable, respectively). Experiments were performed in batch culture with diploid cells using the mother enrichment program (MEP) to enrich aged cells, a well-characterised system that recapitulates lifespan effects of short- and long-lived mutants (47). Cell wall biotin labelling was used to purify aged mother cells, with density gradient separation to remove dead cells and debris (48,49). Cells carried Tom70-GFP as a marker of senescence (6,14), which we have previously shown to be strongly associated with other marks of senescence in the MEP system including vacuolar size, transcriptional dysregulation and chromosomal aberrations (18,50). Cells were co-stained for bud scars using WGA to quantify replicative age (51–53), and an Rpl13a-mCherry control marker that is unaffected by senescence was included in most experiments (54) (Figure 1A).

**Figure 1:**
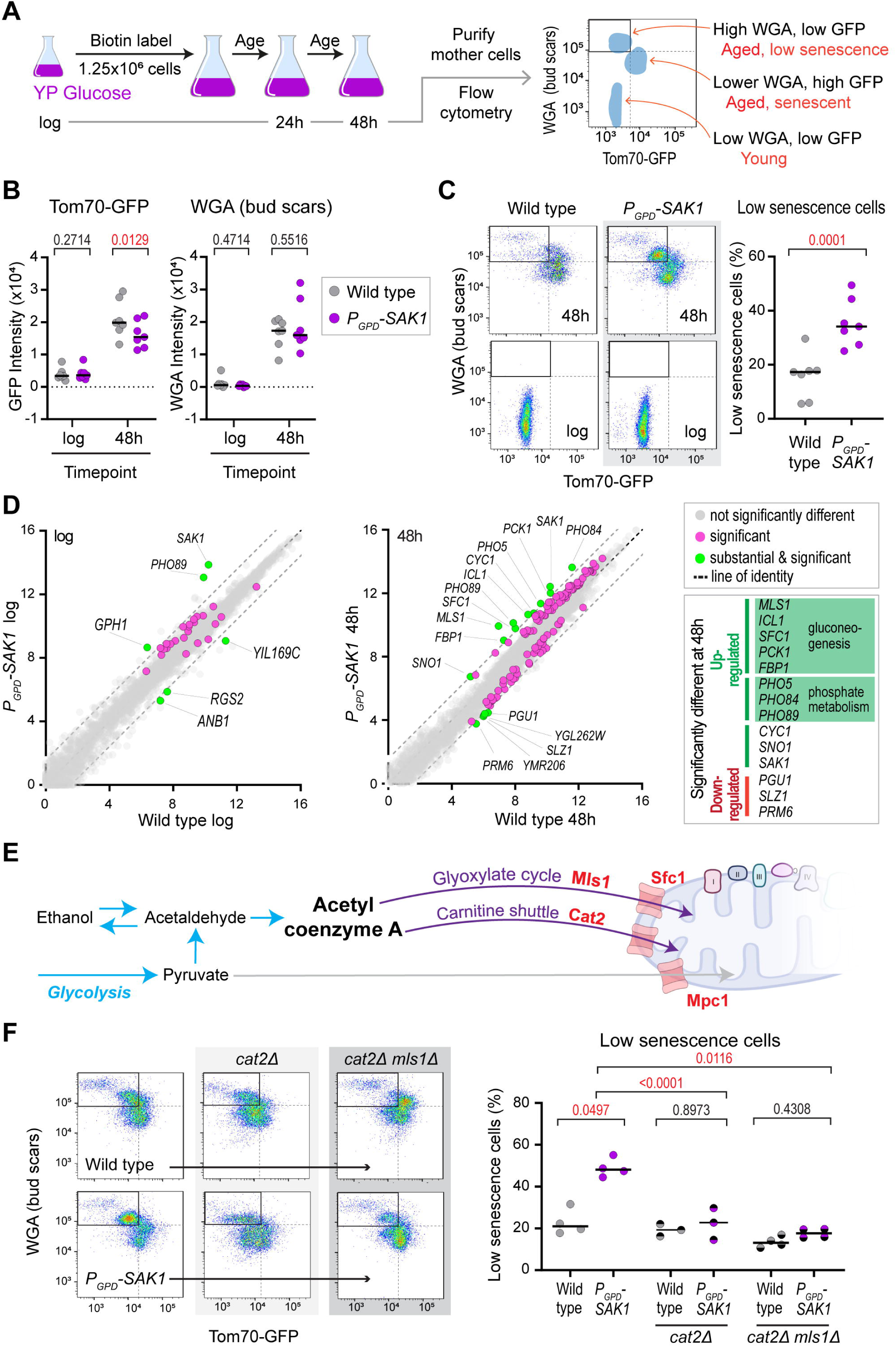
AMPK suppresses senescence through mitochondrial Acetyl-CoA import pathways. **A:** Experimental scheme for analysis of replicatively aged yeast **B:** Impact of *SAK1* overexpression on Tom70-GFP (marker of senescence onset) and WGA (marker of replicative age). p-values calculated by two way ANOVA, n=7 **C:** Separation of wild type and *P_GPD_-SAK1* low senescence (high WGA low Tom70-GFP) and high senescence (low WGA high Tom70-GFP) populations by flow cytometry, and quantification of low senescence population at 48 h. p-value calculated by t test, n=7 **D:** Comparison of gene expression profiles of wild type and *P_GPD_-SAK1* cells at log phase and after 48 h ageing by mRNA-seq. Data is an average of 3 biological replicates (log) or 4 biological replicates (48 h), significantly different genes (p<0.01 by DEseq2) are shown in purple, genes both substantially (>3x indicated by dotted lines) and significantly different are highlighted in green. Substantially and significantly upregulated genes are annotated. **E:** Simple schematic of the relationship between AMPK, mitochondria and glycolysis pathways. Central glucose processing by glycolysis leading to fermentation is shown by blue arrows, AMPK activated pathways as purple arrows. Key enzymes in this study are shown in red. **F:** Representative flow plots and quantification of the low senescence population at 48 h in wild type and *P_GPD_-SAK1* mutants lacking the carnitine shuttle (*cat2*Δ) and mutants lacking both the glyoxylate cycle (*mls1*Δ) and *cat2*Δ. p-values calculated by one way ANOVA, n=3-4

AMPK was activated on glucose by constitutive overexpression of the upstream kinase Sak1 using a *P_GPD_* promoter as previously described (55,56) (Figure S1A). After 48 h ageing, only a subtle decrease in the Tom70-GFP senescence mark was detected in *P_GPD_*-*SAK1* cells by bulk measurements (Figures 1B, S1B), but a new subpopulation of aged cells with low Tom70-GFP and high WGA emerged (Figures 1A, 1C top left quadrant). This subpopulation has a greater replicative age (marked by WGA) since non-senescent cells continue to cycle at normal speed rather than slowing down after the SEP, and so reach a higher replicative age in 48 h. Low Tom70-GFP and high replicative age therefore mark a sub-population of aged cells with reduced senescence that is specific to the *P_GPD_*-*SAK1* mutant (Figure 1C right).

To understand how *P_GPD_*-*SAK1* reduces senescence, we performed RNA-seq comparing *P_GPD_-SAK1* to wild type cells at log and 48 h. Although AMPK is a major regulator of gene expression (57), only 21 genes were significantly differentially expressed at log phase in *P_GPD_-SAK1* (Figure 1D, left), but a greater effect was observed in aged *P_GPD_-SAK1* cells: 121 genes were significantly different from wild type (p<0.01) amongst which the up-regulated genes were enriched for respiration and cytoplasmic translation functions, while the down-regulated genes were not enriched for any GO terms. However, only 16 of these genes were substantially different (>3-fold: 5 down, 11 up including *SAK1*, Figure 1D right): the down-regulated genes shared no apparent connection, but the 11 up-regulated genes included all 5 of a co-regulated gene set that directs cytosolic Acetyl-CoA into gluconeogenesis via the glyoxylate cycle - *MLS1*, *ICL1*, *PCK1, SFC1* and *FBP1* (58,59). This pathway is not used during normal growth on glucose, though plentiful cytosolic Acetyl-CoA is available, but it is required during growth on carbon sources such as ethanol and acetate when Acetyl-CoA becomes the fuel for respiration as well as the feedstock for gluconeogenesis and associated anabolic pathways.

The glyoxylate cycle converts cytosolic or peroxisomal Acetyl-CoA into succinate for transport into mitochondria by Sfc1. Deletion of *MLS1* or *SFC1* significantly reduced the low senescence sub-population in *P_GPD_-SAK1* (Figure S1C), however Acetyl-CoA can also be transported from the cytoplasm into mitochondria after conjugation to carnitine by carnitine acetyl transferases (Figure 1E). Although multiple carnitine acetyl transferases exist outside mitochondria, this carnitine shuttle depends on the mitochondrial carnitine acetyltransferase Cat2 and deletion of *CAT2*, either alone or in combination with *mls1*Δ almost completely abrogated the effect of *P_GPD_-SAK1*, with very few cells achieving a low senescence phenotype at 48 h (Figure 1F).

The transport of cytosolic Acetyl-CoA into mitochondria fuels respiration during growth on acetate and ethanol, and our results indicate that this process also fuels respiration during ageing on glucose. Indeed, the low senescence population in *P_GPD_-SAK1* is dependent on the electron transport chain and does not form in *P_GPD_-SAK1 cox9*Δ cells, but is independent of pyruvate derived from glycolysis entering mitochondria directly via the mitochondrial pyruvate carrier Mpc1 (Figure S1D). Therefore, AMPK activation on glucose maintains mitochondrial activity and prevents the decline of mitochondria marked by high Tom70-GFP intensity, which was previously described as the cause of senescence on one yeast ageing trajectory (6,60).

### Combining AMPK and Acc1 activity to prevent senescence

AMPK only suppresses senescence in a sub-population of cells, defining two classes of ageing cells – those in which senescence is reduced by AMPK activation and those in which AMPK activation is ineffective. This led us to ask whether AMPK activation also has a negative impact that dominates in the latter class. Having implicated Acetyl-CoA in senescence, we investigated other fates of this metabolite that are also regulated by AMPK. Cytosolic Acetyl-CoA is the building block for fatty acids, being processed into malonyl coenzyme A by ACC to initiate fatty acid synthesis, however ACC is inhibited by AMPK to suspend energy storage in fatty acids under low energy conditions. Yeast ACC (Acc1) is inhibited by AMPK through phosphorylation at two serines, with Ser1157 being the main regulatory site (Figure 2A) (61,62). To test whether Acc1 inhibition drives senescence in the sub-population of cells that do not respond to Sak1 over-expression, we combined *P_GPD_-SAK1* with an *acc1^S1157A^* mutation that prevents Acc1 inactivation by AMPK, thereby creating the A2A mutant (<u>A</u>MPK and <u>A</u>cc1 <u>a</u>ctive).

**Figure 2:**
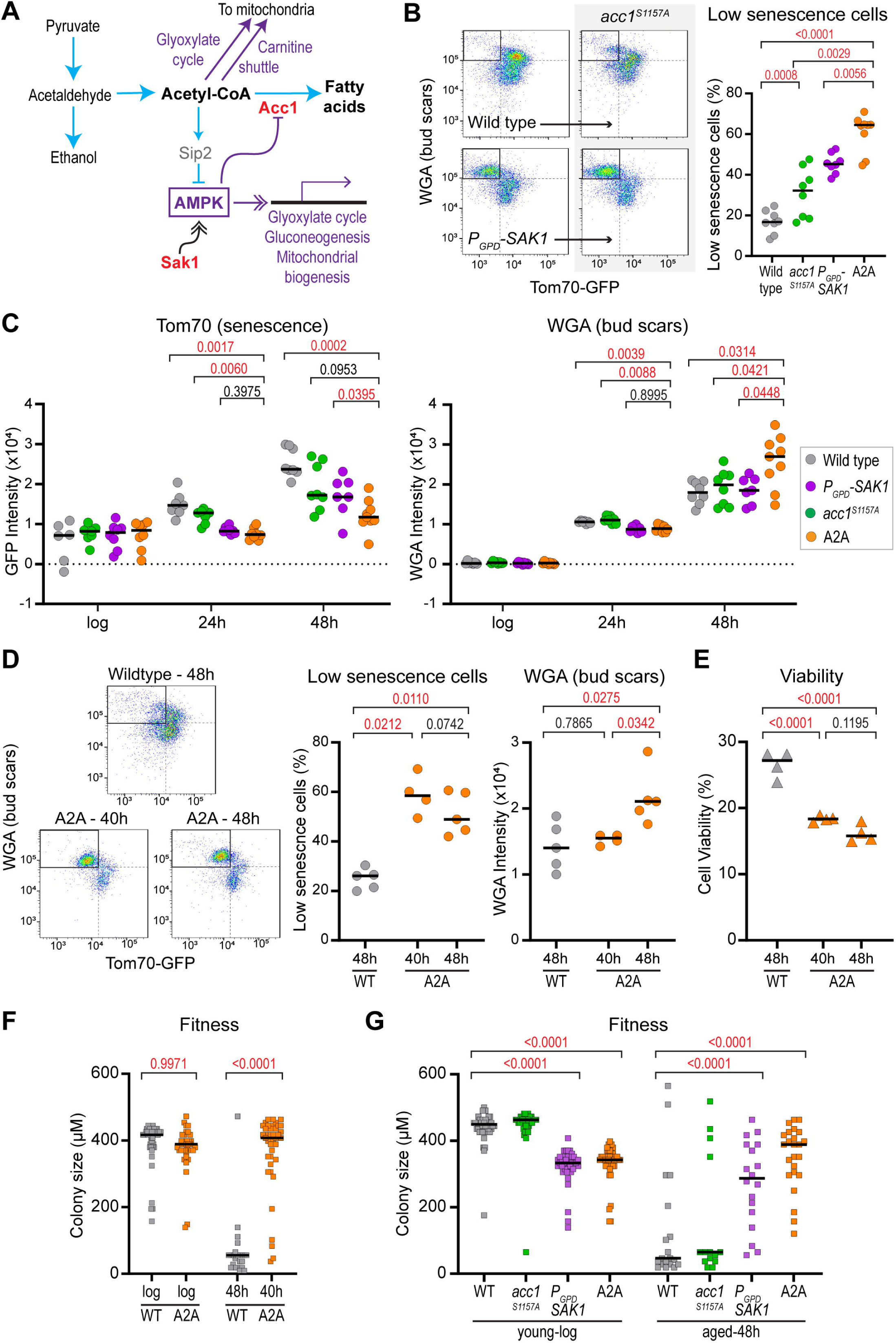
Combining AMPK activity and fatty acid synthesis to avoid senescence. **A:** Schematic highlighting the regulation of fatty acid synthesis by AMPK. Key proteins for this figure are highlighted in red. **B:** Representative flow plots and quantification of the low senescence population in acc1^S1157A^, *P_GPD_-SAK1* and A2A at 48 h. p values calculated by one way ANOVA, n=8-9 **C:** Population medians for Tom70-GFP and WGA in acc1^S1157A^, *P_GPD_-SAK1* and A2A at log phase, 24 h and 48 h. p values calculated by two way ANOVA, n=7-9 **D:** Representative flow plots of wild type and A2A at matched average replicative age (48 h for wild type versus 40 h for A2A), with quantification of the low senescence population and of median WGA (bud scars). p values calculated by one way ANOVA, n=4-5 **E:** Percentage cell viability in cultures aged for 40 and 48 h. p value calculated by t test, n=4. **F:** Size of colonies formed in 24 h on a YPD plate by log phase and age-matched wild type and A2A. Only cells that eventually formed colonies within 3 days were included to ensure all tested cells were viable. p values calculated by two way ANOVA, n=14-39 **G:** Size of colonies formed in 24 h on a YPD plate by log phase and 48 h-aged wild type, acc1^S1157A^, *P_GPD_-SAK1* and A2A. Only cells that eventually formed colonies within 3 days were included to ensure all tested cells were viable. p values calculated by two way ANOVA, n=15-61

Young A2A cells proliferate equivalently to wild-type cells in liquid culture (Figure S2A), and do not have an extended replicative lifespan (Figure S2B), but do show a significant and substantial increase in the low senescence population relative to *P_GPD_-SAK1* (Figure 2B). Separate analysis of Tom70-GFP and WGA revealed that the A2A and *P_GPD_-SAK1* populations were equivalent on average at 24 hours, but A2A consistently presented lower Tom70-GFP and higher WGA by 48 h (Figures 2C, S2C, S2D). Population measurements of Tom70-GFP and WGA do vary between experiments, but the differences were consistent across many experiments (Figure S2D, n>25), and bud scar counting confirmed that A2A cells reach a higher age than wild-types (Figure S2E). The combination of AMPK activation and unfettered Acc1 activity therefore allows the majority of cells to age without senescence. A2A cells cultured for 40 h reached an average replicative age comparable to wild-type at 48 h and, at this point, formed a very clear low senescence population, confirming that SEP onset is delayed or absent in age-matched cells (Figure 2D). This reduction of senescence occurred despite the viability of A2A cells (based on the ability to form a colony on a YPD plate) being lower at 40 h than wildtype at 48 h (Figure 2E). Importantly, at 40 h, >50% of the A2A cells are in the low Tom70-GFP, high age population even though <20% are viable, meaning that many cells reach the end of their replicative lifespan without passing through the SEP.

Tom70-GFP and WGA signals are only indirect indicators of cell fitness, whereas the growth rate of colonies formed by individual young or old cells provides a direct physiological measure of cell fitness. While young A2A cells formed slightly smaller colonies than did wild-type cells, the colonies formed by aged A2A cells were much larger than those formed by wild-type cells of equivalent replicative age (Figure 2F). Indeed, aged A2A cells formed colonies of similar size to young A2A cells, showing equivalent fitness, whereas most aged wild-type cells divided only once or not at all across the 24-hour assay period. Only viable cells that formed colonies within 3 days were included in this analysis, so the non-dividing aged wild-type cells were viable but had a cell division time greater than 24 hours. This demonstrates that A2A cells maintain excellent fitness late in life, whereas the fitness of aged wild-type cells is very low. A comparison of the fitness of the wild type, *acc1^S1157A^*, *P_GPD_-SAK1* and A2A strains at a fixed 48 h time point revealed that A2A maintained the best fitness despite also having the highest median age, while *P_GPD_-SAK1* showed variable fitness, and *acc1*^S1157A^ was similar to wild type (Figure 2G). It is worth noting that colony formation by log phase cells carrying the *P_GPD_-SAK1* construct was consistently slightly slower than wildtype (5% longer doubling time on average) although this does not manifest in liquid growth assays.

Senescent onset in haploid yeast has been linked to the accumulation of extrachromosomal ribosomal DNA circle (ERC) (63,64), but ERCs accumulate to a greater extent in A2A than in wildtype (Figure S2F) consistent with our previous finding that ERCs are not linked to senescence in diploids (50). SEP onset in diploids is associated with accumulation of a fragment of chromosome XII (ChrXIIr) detectable through increased expression of genes in this ∼600kb genomic region (50); ChrXIIr accumulation is reduced in *P_GPD_-SAK1*, but surprisingly not further reduced in A2A (Figure S2G), indicating that ChrXIIr formation is not sufficient to cause senescence and that preventing Acc1 inhibition rescues senescence by a mechanism that does not reduce ChrXIIr formation.

These experiments show that the beneficial effects of AMPK activation during ageing can be increased substantially if Acc1 inhibition is prevented, and are independent of previously reported genome instability phenotypes associated with the rDNA. High AMPK activity combined with normal Acc1 activity results in a robust low senescence ageing phenotype and high late life fitness without decreasing mortality rate.

### Different classes of A2A cells depend on respiration and Acc1 activity

We then asked whether all A2A cells avoid senescence through the same mechanism. Given that the required factors (Acc1, the carnitine shuttle, the glyoxylate cycle) each deplete the cytosolic Acetyl-CoA pool, a prime candidate is the negative feedback loop involving the AMPK subunit Sip2, which inhibits AMPK when cytosolic Acetyl-CoA availability is high (65) (Figure 3A). It is not hard to imagine that the metabolic balance emerging from this negative feedback loop varies between cells, with some cells being more dependent than others on cytosolic Acetyl-CoA removal by Acc1 to maintain AMPK activity. Depleting Acetyl-CoA reduces bulk histone acetylation (66), and we observed this effect in A2A cells, indicating that cytosolic -CoA availability is indeed decreased (Figure 3B). However, AMPK activation with *P_GPD_-SAK1* yielded the same split population in the *sip2*Δ strain as in wildtype (compare Figs. 1C, 3C), and the progressive increase in the low senescence population achieved with *P_GPD_-SAK1* and *P_GPD_-SAK1 acc1^S1157A^* (A2A) was also observed in a *sip2^3R^* mutant that does not respond to Acetyl-CoA (65)(Figure S3A). Therefore, the differential responses to AMPK activation in the ageing population cannot be attributed to Sip2-mediated feedback, but depletion of cytosolic Acetyl-CoA may be beneficial independent of Sip2.

**Figure 3:**
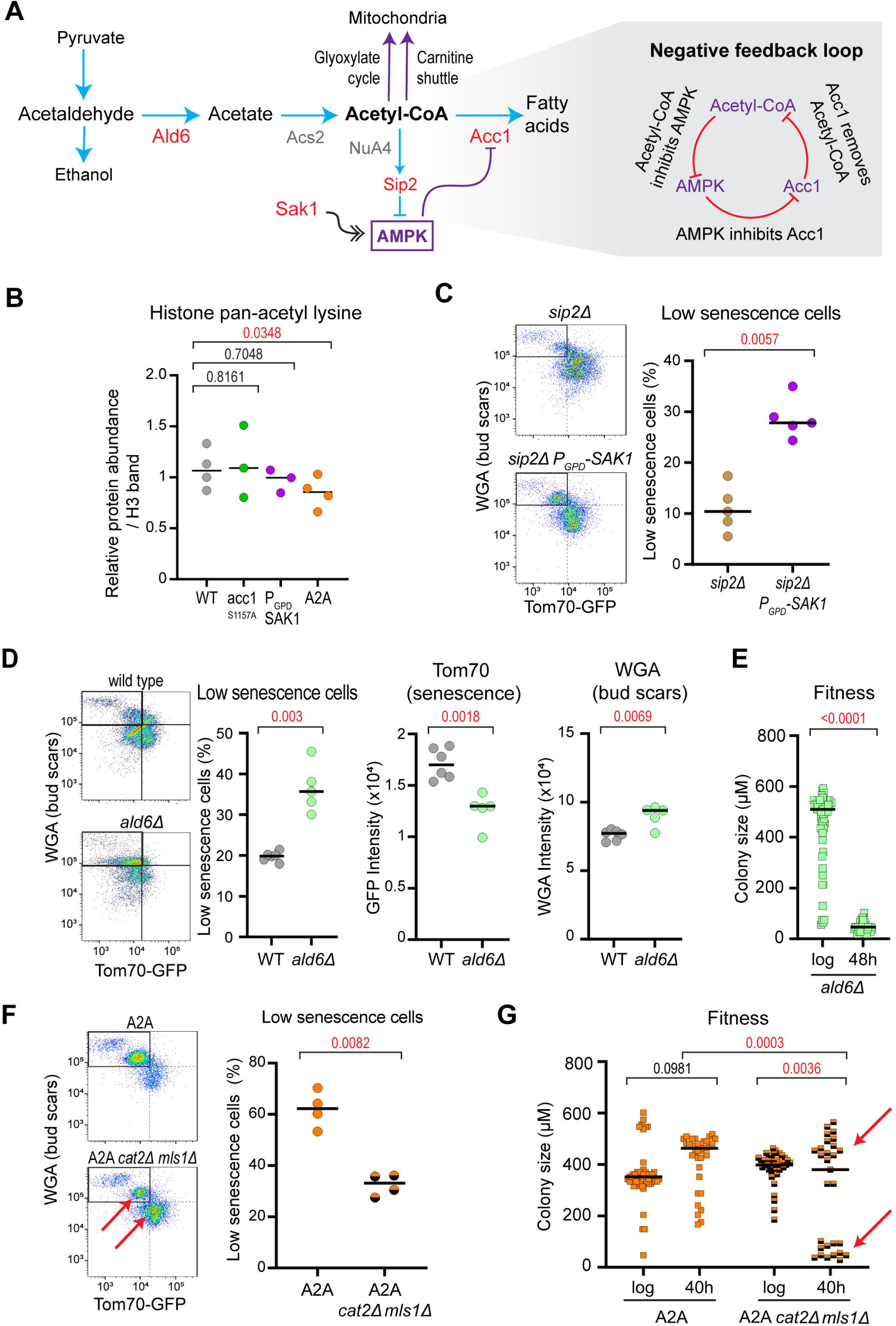
Separate classes of A2A cells depend on respiration and fatty acid availability. **A:** Schematic of acetyl coenzyme A biosynthesis and the emergence of a negative feedback loop through AMPK regulation by Sip2. Key proteins for this figure are highlighted in red. **B:** Western blot analysis of acetyl lysine on H3 in log phase wild type, *acc1^S1157A^*, *P_GPD_-SAK1* and A2A cells. Membranes were probed with mouse anti-H3 and rabbit anti-pan-acetyl lysine using a 2-colour detection system. p values calculated by one-way ANOVA, n=3-4. **C:** Representative flow plots and quantification of low senescence population in *sip2*Δ and *sip2*Δ *P_GPD_-SAK1* at 48 h. p-values calculated by t-test, n=5 **D:** Representative flow plots and quantification of low senescence population in wild type and *ald6*Δ at 48 h. Population medians are also shown for Tom70-GFP and WGA in wild type and *ald6*Δ at 48 h. p values calculated by t-test, n=5-6 **E:** Size of colonies formed in 24 h on a YPD plate by log phase and 48 h-aged wild type and *ald6*Δ. Only cells that eventually formed colonies within 3 days were included to ensure all tested cells were viable. p values calculated by one-way ANOVA, n=36-57 **F:** Representative flow plots and quantification of low senescence population in A2A mutants lacking both carnitine acetyltransferase activity (*cat2*Δ) and the glyoxylate cycle (*mls1*Δ) at 48 h. Arrows indicate two populations, p-value calculated by t-test, n=4 **G:** Size of colonies formed in 24 h on a YPD plate by log phase and 40 h-aged A2A and A2A *cat2*Δ *mls1*Δ. Only cells that eventually formed colonies within 3 days were included to ensure all tested cells were viable. Arrows indicate two populations, p values calculated by two way ANOVA, n=29-55

To determine whether reducing the availability of Acetyl-CoA is beneficial in ageing, we deleted *ALD6*, which encodes the primary cytoplasmic acetaldehyde reductase. Loss of Ald6 reduces the supply of acetate from fermentation to be processed into Acetyl-CoA (67,68). Although this did not have a large effect on the low Tom70-GFP, high WGA population as defined in Figure 1, the deletion of *ALD6* was effective in decreasing Tom70-GFP and increasing age at harvest (Figure 3D). These findings showed that *ald6*Δ cells maintain high fitness for a longer portion of life. However, fitness measured at 48 h was very low (Figure 3E) meaning that the fitness gains from reducing cytosolic Acetyl-CoA availability are temporary and are lost by 48 h; therefore, the A2A phenotype cannot be attributed solely to a reduction of Acetyl-CoA availability.

Conversely, the low senescence of A2A populations may have different causes in different cells, with some cells requiring AMPK-driven respiration fuelled by cytosolic Acetyl-CoA and other cells requiring the combination of high AMPK and high Acc1 activity. To test this hypothesis, we deleted C*AT2* and *MLS1* in A2A, which should impair ageing fitness of only those cells requiring AMPK-driven respiration: this resulted in the population dividing between low and high senescence states based on flow cytometry and fitness (Figure 3F,G). Similarly, deleting *COX9* to prevent respiration removed a subset of cells from the low senescence population (Figure S3B). Therefore, cells that achieve a low senescence state through respiration in *P_GPD_-SAK1* avoid senescence by the same mechanism in A2A, but the cells that avoid senescence due to the *acc1^S1157A^* mutation in A2A use a mechanism independent of the carnitine shuttle and glyoxylate cycle. We further asked whether respiration itself or reduction in Acetyl-CoA availability is the primary cause of senescence reduction for the first group of cells by deleting *ALD6* in *P_GPD_-SAK1*; if respiration is the primary cause, *ALD6* deletion should suppress the benefit of *P_GPD_-SAK1* by reducing the fuel for respiration, whereas if Acetyl-CoA removal is the primary cause then this should be neutral or beneficial. Working in *sip2*Δ to avoid an increase in AMPK activity due to reduced Acetyl-CoA availability, we observed that *ald6*Δ increased the low senescence population through decreasing Tom70-GFP (S3C), and therefore the beneficial impact of *P_GPD_-SAK1* on this pathway arises primarily through Acetyl-CoA removal. It is possible that respiration is adding to this benefit, and we detect a significant increase in Oxygen Consumption Rate in aged *P_GPD_-SAK1* cells, but the further increase in A2A is smaller and we consider that this cannot fully explain the rescue achieved by *acc1^S1157A^*.

Cytosolic Acetyl-CoA is a key regulator of histone acetylation and thereby chromatin accessibility and gene silencing, so increased acetyl-CoA availability could affect gene expression in multiple ways. Dysregulation of gene expression with age, or more exactly with senescence, causes induction of genes not normally expressed during growth in glucose (18,50,52,69–71). AMPK promotes the transcription of many of these genes (57), but age-linked transcriptional dysregulation is reduced in *P_GPD_-SAK1* compared to wildtype indicating better maintenance of a youthful gene expression profile (Figure S3E). However, gene expression dysregulation in A2A is equivalent to *P_GPD_-SAK1* and gene expression patterns are almost identical with no significantly differentially mRNAs (Figure S3E,F). We have previously noted that Sir2-repressed genes (primarily from sub-telomeric regions) are not dysregulated with age to a greater extent than other genes (50), and although transcripts from the Sir2-repressed rDNA intergenic spacer regions are massively induced with age, we attribute this to the increase in copy number of these loci through ERC accumulation (Figures S2F,S3E). These species increase less with age in A2A, but this reflects higher abundance at log phase rather than lower abundance in aged cells. In summary, a proportion of age-linked gene expression dysregulation may be linked to cytosolic acetyl-CoA availability, but our evidence does not indicate a major effect on HDAC-mediated silencing.

These experiments define two classes of ageing cells with distinct metabolic needs, coherent with the model of two ageing trajectories previously proposed (6). In one class, respiration fuelled by cytosolic Acetyl-CoA is required to suppress senescence, while in the other class respiration is not required but maintaining Acc1 activity is critical. These processes all consume cytosolic Acetyl-CoA, and reducing the availability of this metabolite is beneficial but insufficient to prevent senescence.

### Fatty acid supply is limiting in ageing cells that do not respond to AMPK activation

Acc1 converts Acetyl-CoA to Malonyl-CoA as the first step in fatty acid synthesis, a biosynthetic activity that may or may not be important during ageing. We asked whether fatty acids or perhaps the Malonyl-CoA intermediate prevent senescence onset in A2A, against a null hypothesis that Acc1 simply provides an alternative pathway to respiration for moderating cytosolic Acetyl-CoA levels (Figure 4A). To test this hypothesis, we attempted to reproduce the low senescence phenotype of A2A by manipulating fatty acid synthesis in *P_GPD_-SAK1,* again in the absence of Sip2 to avoid feedback on AMPK activity (Figure 4B).

**Figure 4:**
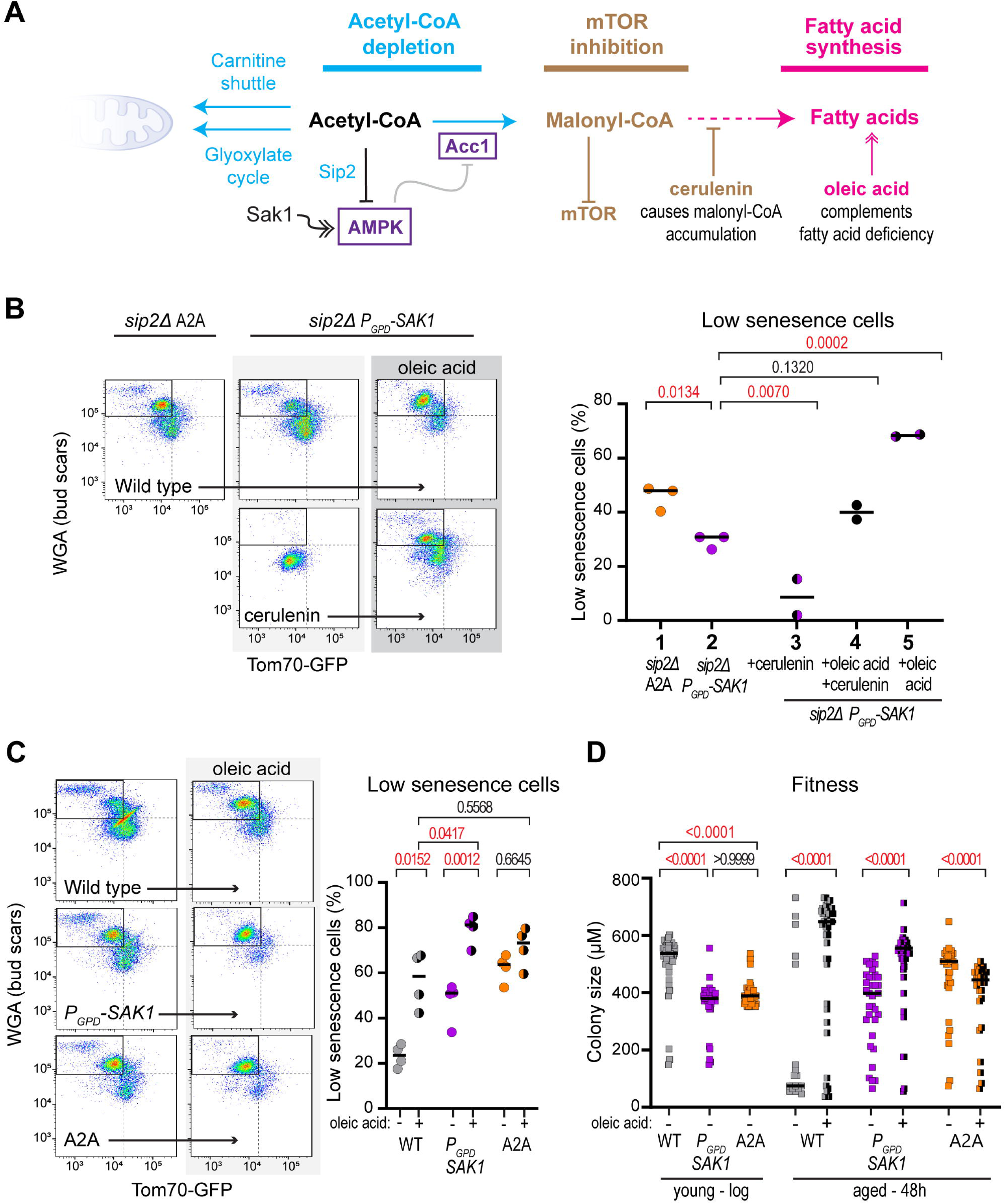
Fatty acids are particularly important in ageing cells that do not respond to AMPK activation. **A:** Schematic of potential mechanisms for the beneficial effect of ACC1 activity: Inhibition of mTOR mediated by malonyl coenzyme A, the direct product of Acc1, or efficient synthesis of fatty acids, compared to a null hypothesis that Acc1 acts to remove cytosolic acetyl-CoA, for example if these cells are respiratory-deficient. **B:** Representative flow plots and quantification of low senescence population at 48 h in *sip2*Δ *P_GPD_-SAK1 acc1^S1157A^*andL*sip2*Δ *P_GPD_-SAK1* supplemented with 2.5 µM cerulenin (condition 3), 10 µM cerulenin + 0.04% oleic acid + 0.04% Tween 80 (condition 4) or 0.04% oleic acid + 0.04% Tween 80 (condition 5). p-values calculated by one-way ANOVA, n=2-3 **C:** Representative flow plots and quantification of low senescence population at 48 h in wild type, *P_GPD_-SAK1* and A2A supplemented with 0.04% oleic acid + 0.04% Tween 80. p-values calculated by one-way ANOVA, n=4 **D:** Size of colonies formed in 24 h on a YPD plate by log phase and 48 h-aged wild type, acc1^S1157A^, *P_GPD_-SAK1* and A2A supplemented with 0.04% oleic acid + 0.04% Tween 80. Only cells that eventually formed colonies within 3 days were included to ensure all tested cells were viable. p values calculated by one-way ANOVA, n=19-52

Malonyl-CoA was recently found to be an endogenous mTOR inhibitor (72), and mTOR inhibition is well known to increase ageing health and lifespan (11). However, a low dose of the fatty acid synthase inhibitor cerulenin, which causes Malonyl-CoA to accumulate (72,73), profoundly decreased the low senescence population (Figure 4B, conditions 2,3). A higher dose of cerulenin was also tested in combination with oleic acid supplementation to complement the complete blockade of fatty acid synthesis, but this combination had very little effect (Figure 4B, condition 4). Therefore, Acc1 activation in A2A cells does not work through increased Malonyl-CoA levels. In contrast, supplementation with only the monounsaturated fatty acid oleic acid increased the low senescence population of *P_GPD_-SAK1* cells even more than *acc1^S1157A^*, showing that fatty acid scarcity is the primary driver of senescence in ageing cells that do not respond to AMPK activation alone.

None of these treatments had a strong impact on Tom70-GFP however, with increases in the low Tom70-GFP, high WGA population being driven almost entirely by increased WGA (Figure S4A), indicating that the strongest effect of fatty acid supplementation was to overcome the known short lifespan of the *sip2*Δ background (37). After validating the effect of oleic acid by bud scar counting (Figure S4B), we therefore asked whether oleic acid supplementation also reduces senescence in a wild-type background. The effects were dramatic, with the low senescence population increasing 2.5-fold through decreased population average Tom70-GFP and increased WGA (Figures 4C, S4C). The low senescence population in *P_GPD_-SAK1* also almost doubled with oleic acid supplementation, but only a small, non-significant effect on A2A was observed (Figure 4C). Notably, oleic acid supplementation and *P_GPD_-SAK1* each allow approximately half of the population to age with low senescence, while the combination allows most cells to avoid senescence - this is as expected if *P_GPD_-SAK1* and oleic acid supplementation suppress senescence in different classes of ageing cells.

At 48 h the fitness of wildtype cells supplemented with oleic acid also increased dramatically, *P_GPD_-SAK1* cells improved somewhat while fitness was significantly but not substantially decreased in A2A (Figure 4D). In fact, the oleic acid-supplemented wild-type cells were fitter than the mutants, which likely reflects the decrease in baseline fitness caused by *P_GPD_-SAK1* in young cells as noted above. The results for *P_GPD_-SAK1* confirmed that a subset of cells in which AMPK is activated still undergo senescence because of fatty acid starvation, which can be compensated by the *acc1^S1157A^* mutation or by oleic acid supplementation.

However, the results for the wild type are more complicated as ∼40% of oleic acid-supplemented cells undergo senescence according to flow cytometry whereas all oleic acid supplemented wild-type cells are extremely fit (compare Figures 4C and 4D). This difference arises because fitness assays exclude inviable cells, such as the cells that senesce and lose viability before 48 h due to lack of respiration, which are exactly the cells for which senescence is rescued by AMPK activation. We further noted that the low fitness of aged *ald6*Δ cells was rescued by oleic acid to wild-type levels, showing that the observed fitness defect simply arose from fatty acid starvation rather than from any other function of Acetyl-CoA (Figure S4D). In keeping with observations in A2A, oleic acid suppresses senescence and restores fitness in a large fraction of the population without reducing average age-linked gene expression dysregulation or ChrXIIr accumulation (Figure S4E-G), indicating that neither phenotype is sufficient to cause senescence. 115 genes are significantly different in cells aged on oleic acid, most of which are upregulated and are highly enriched for translation reflecting the increased fitness and rapid growth, and it is the higher expression of these ribosomal protein genes that reduces the negative slope value that quantifies gene expression dysregulation compared to untreated wild types (compare Figures S3E, S4E).

Overall, in wild-type cells, as in *P_GPD_-SAK1 cells*, high Acc1 activity or oleic acid supplementation prevents senescence in one class of ageing cells while AMPK activation prevents senescence in the other class.

## Discussion

Our findings indicate a major role for Acetyl-CoA metabolism in the yeast replicative ageing process. Combining transport of cytosolic Acetyl-CoA into mitochondria with ongoing cytosolic fatty acid synthesis together ensures that cells rarely enter senescence. This is achieved without reducing age-linked mortality and with little effect on the fitness of cells when young, showing that lifespan and fitness decline are separable in the yeast replicative ageing system, and that high fitness can be maintained to the end of life – an effect known as compression of morbidity.

Here we have characterised two classes of ageing cells seemingly differentiated by high and low availability of cytosolic Acetyl-CoA, consistent with a previous demonstration that ageing follows two trajectories in yeast though it should be noted that in this previous report, the two trajectories also differed in replicative lifespan (6). Acetyl-CoA is the feedstock for fatty acid synthesis, which is necessary both for membrane production and for energy storage in lipid droplets. However, cytosolic Acetyl-CoA production consumes as much ATP as is generated in glycolysis and thus competes directly with growth. The existence of two classes of cell that follow different ageing trajectories makes sense because yeast cells do not know when food availability will change and so the population employs a bet hedging strategy (74–76): part of the population synthesises excess cytosolic Acetyl-CoA to store as much energy as possible in lipids against future scarcity, while other cells maximise growth from the current food supply by synthesising the bare minimum of cytosolic Acetyl-CoA needed for membrane formation (Figure 5). Our findings indicate that though both strategies are useful they eventually decrease fitness as cells age, forming a classic antagonistic pleiotropy (3).

**Figure 5:**
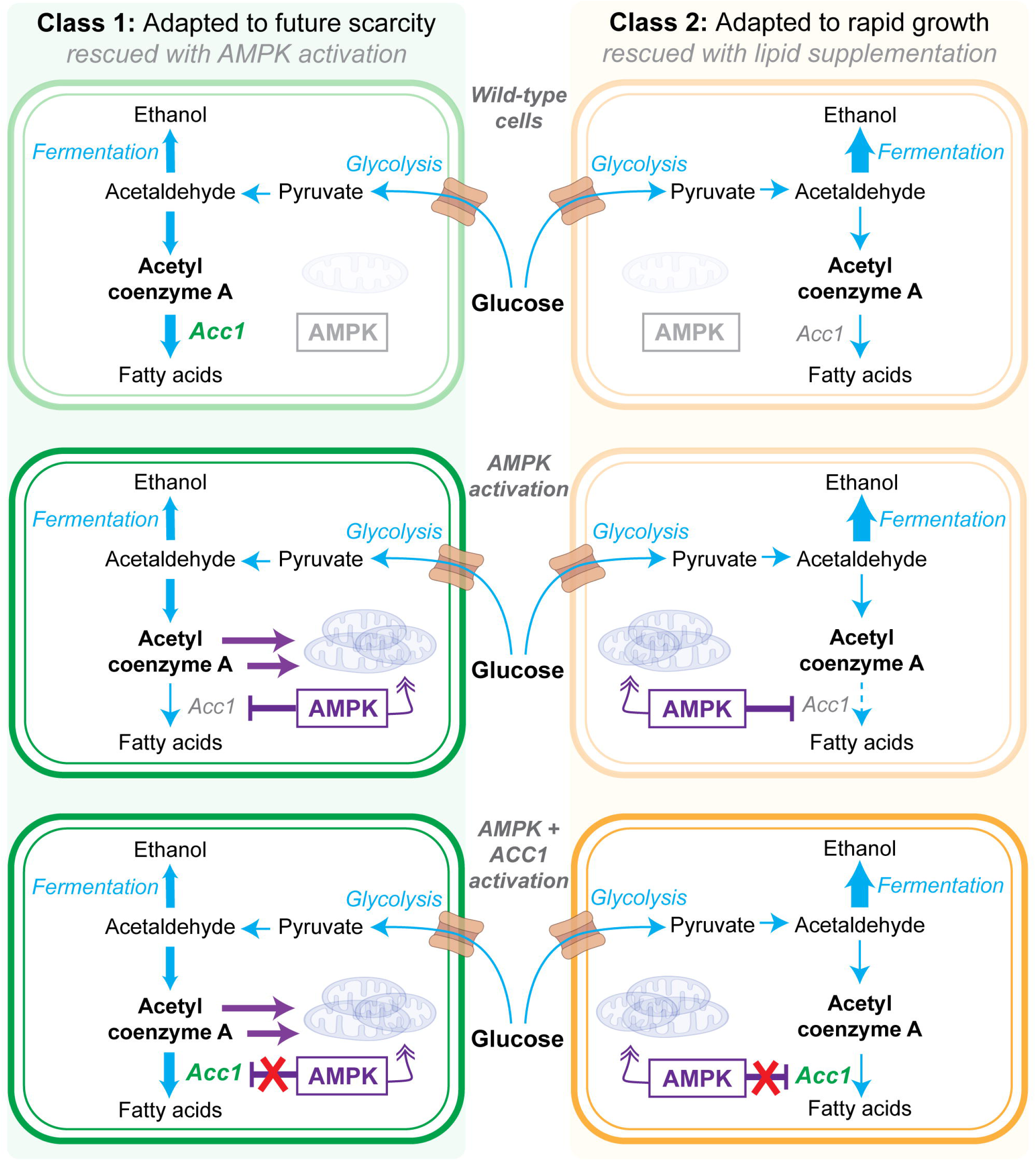
Model of different classes of ageing mediated by Acetyl-CoA and lipid metabolism. Schematic demonstrating the two wild type senescence fates. (i) Adaptation to future scarcity by synthesising excess cytosolic Acetyl-CoA to store as much energy as possible in lipids against future scarcity, and (ii) Adaptation to rapid growth by maximising growth from the current food supply by synthesising the bare minimum of cytosolic Acetyl-CoA.

Separation of ageing trajectories proves useful in identifying ageing phenotypes that may be causal for fitness differences from correlated bystanders. While low Acetyl-CoA transport to mitochondria and lipid synthesis promote senescence in different cells, we find that age-linked gene expression dysregulation can be decoupled from fitness as can accumulation of ChrXIIr. Both phenotypes correlated tightly with the SEP in our previous analyses (18,50), but A2A and oleic acid rescue fitness without rescuing either of these phenotypes, though we cannot exclude that these phenotypes contribute to senescence in cells that are rescued by AMPK activation. Our results also confirm our previous findings that ERC accumulation and loss of chromatin silencing are unrelated to the SEP, at least in diploid cells (50).

In general, multiple pathways can promote senescence during yeast ageing; which pathway dominates is dependent on the experimental system and therefore the set of stresses experienced by cells (6,15,17,77,78). The SEP in yeast was originally characterised in microfluidic assays, where it is attributed to the accumulation of extrachromosomal rDNA circles (ERCs) (14,15,78). In contrast, ERC accumulation is completely unrelated to senescence during ageing in the batch culture system employed here (18,48,50), which is sensitive to metabolic changes such as a galactose diet or Acetyl-CoA processing; the galactose diet has no effect on cells in microfluidic systems and the impact of caloric restriction has remained controversial, so we do not necessarily expect A2A to prevent senescence in microfluidic systems (79,80). However, our findings parallel studies of yeast chronological ageing which have highlighted the importance of respiration and lipid synthesis, though chronological ageing measurements are performed in glucose-depleted conditions and it has been difficult to translate these findings to ageing under normal nutrient availability (23,66,81). Whatever the root of the difference, ageing in batch culture allows the exploration of ageing biology with little impact from ERC accumulation, a phenomenon that does not occur during ageing in animals.

AMPK is broadly considered to have a pro-longevity effect, however the benefits of AMPK activation under normal diets are limited, and some studies in budding yeast conversely found that AMPK activity is negative for lifespan (82,83). Our work shows that both positions can be true, with AMPK having both beneficial and detrimental impacts under nutrient replete conditions. It is not too surprising that issues arise when AMPK is ectopically activated on high glucose given that AMPK is a central energy regulator that evolved to manage metabolism under low nutrient conditions, and it is perhaps more surprising that a single point mutation largely corrects these issues.

The AMPK-Acc1 regulatory system is conserved from yeast to humans, but the source and fate of cytosolic Acetyl-CoA are different so the combination of AMPK activation with lipid supplementation is unlikely to be universally effective. In human cells, the only major route for removal of excess cytosolic Acetyl-CoA is lipid synthesis to palmitoyl coenzyme A followed by carnitine-dependent transport into mitochondria for oxidation. Ectopic activation of AMPK under high nutrient availability would promote lipid catabolism but inhibits lipid synthesis through ACC1/2 phosphorylation. Preventing inhibition of ACC1/2 by AMPK is ineffective as ACC2 also inhibits fatty acid catabolism so fatty acid synthesis and storage increase, resulting in non-alcoholic fatty liver disease (84). However, we speculate that activating AMPK while selectively preventing AMPK inhibition of ACC1 alone may be effective.

## Materials and Methods

Detailed and updated protocols are available at https://www.babraham.ac.uk/our-research/epigenetics/jon-houseley/protocols.

### Yeast strain construction, culture and labelling

Yeast deletion mutants and fusion constructs were constructed by standard methods and are listed in Table S1, oligonucleotide sequences are given in Table S2. Plasmid templates were pFA6a-GFP-KanMX4, pYM-N14, pAW8-mCherry, pFA6a-HIS3 and pFA6a-URA3 (85–87). To create the *acc1^S1157A^* point mutation, *ACC1* was amplified with ACC1 F3/R3 and cloned in pRS314, then the S1157A mutation introduced by site directed mutagenesis. MEP **<u>a</u>** ±Kan-*P_GPD_-SAK1* cells carrying this plasmid were transformed with pGSKU (88) amplified with ACC1 CORE UP45 1 and ACC1 CORE DN45 2 to replace the chromosomal ACC1 S1157 codon with a CORE cassette flanked on each side by I-*Sce*I sites. Cells were plated on YPGal to induce I-*Sce*I then FOA to select for loss of the CORE cassette, then TRP negative cells lacking the pRS314-*acc1^S1157A^* plasmid were verified by sequencing. *sip2^3R^* was constructed by a simpler method as *SIP2* is not an essential gene: a CORE cassette flanked by I-*Sce*I sites was amplified from pGSKU using SIP2 3Q-R CORE UP45 and SIP2 3Q-R CORE DN45 v2 and transformed in destination strains. These were transformed with pRS413 containing an *Eag*I-*Sac*I fragment of *SIP2* with the *sip2^3R^* sequences, then cells plated on YPGal then FOA.

All cells were grown in YPD media (2% peptone, 1% yeast extract, 2% glucose) at 30°C with shaking at 200 rpm. Media components were purchased from Formedium and media was sterilised by filtration. MEP experiments were performed as described with modifications 89: cells were inoculated in 4 ml YPD and grown for 6-8 hours then diluted in 25 ml YPD and grown for 16-18 hours to 0.2-0.6×10^7^ cells/ml in 50 ml Erlenmeyer flasks. 0.125×10^7^ cells per aged culture were harvested by centrifugation (15 s at 13,000 g), washed twice with 125 µl PBS and re-suspended in 125 µl of PBS containing ∼3 mg/ml Biotin-NHS (Pierce 10538723). Cells were incubated for 30 min on a wheel at room temperature, washed once with 125 µl PBS and re-suspended in 125 µl YPD then inoculated in 125 ml YPD at 1×10^4^ cells/ml in a 250 ml Erlenmeyer flask (FisherBrand FB33132) sealed with Parafilm. 1 µM β-estradiol (from stock of 1 mM Sigma E2758 in ethanol) was added after 1.5h. An additional 0.125×10^7^ cells were harvested from the log phase culture while biotin labelling reactions were incubating at room temperature. Cells were harvested by centrifugation for 1 min, 4600 rpm, immediately fixed by resuspension in 70% ethanol and stored at -80°C. To minimise fluorophore bleaching in culture, the window of the incubator was covered with aluminium foil, lights on the laminar flow hood were not used during labelling and tubes were covered with aluminium foil during biotin incubation. Lifespan assays in the MEP background were performed as previously described (47,89).

### Cell purification

Percoll gradients (1-2 per sample depending on harvest density) were formed by vortexing 1 ml Percoll (Sigma P1644) with 110 µl 10x PBS in 2 ml tubes and centrifuging 15 min at 15,000 g, 4 °C. Ethanol fixed cells were defrosted and washed once with 1 volume of cold PBSE (PBS + 2 mM EDTA) before resuspension in ∼100 µl cold PBSE per gradient and layering on top of the pre-formed gradients. Gradients were centrifuged for 4 min at 2,000 g, 4 °C, then the upper phase and brown layer of cell debris removed and discarded. 1 ml PBSE was added, mixed by inversion and centrifuged 1 min at 2,000 g, 4 °C to pellet the cells, which were then re-suspended in 1 ml PBSE per time point (re-uniting samples where split across two gradients). 25 µl Streptavidin microbeads (Miltenyi Biotech 1010007) were added and cells incubated for 5 min on a wheel at room temperature. Meanwhile, 1 LS column per sample (Miltenyi Biotech 1050236) was loaded on a QuadroMACS magnet and equilibrated with cold PBSE in 4 °C room. Cells were loaded on columns and allowed to flow through under gravity, washed with 1 ml cold PBSE and eluted with 1 ml PBSE using plunger. Cells were re-loaded on the same columns after re-equilibration with ∼500 µl PBSE, washed and re-eluted, and this process repeated for a total of three successive purifications. After addition of Triton X-100 to 0.01% to aid pelleting, cells were split into 2 fractions in 1.5 ml tubes, pelleted 30 s at 20,000 g, 4 °C, frozen on N2 and stored at -70 °C.

### Fitness and viability assays

For live purification, cells were pelleted, washed twice with synthetic complete 2% glucose media, then resuspended in 2 ml final volume of the same media and incubated for 5 min on a rotating wheel with 10 µl MyOne streptavidin magnetic beads (ThermoFisher), isolated using a magnet and washed five times with 1 ml of the same media. Cells were streaked on a YPD plate and individual cells moved to specific locations using a Singer MSM400 micromanipulator. Colony size was measured 24 hours later on the screen of the micromanipulator imaging with a 4x objective, sizes were calibrated using a haemocytometer. Cells that did not form a colony within 72 h were assumed to have been inviable and were not included in the analysis. Colony growth speed is defined by daughter cell replication, and if daughters and subsequent generations divide at the same rate irrespective of whether they come from young or old mothers then the size of the colony after 24 hours varies simply based on the time it took the initial mother to produce a daughter (12). In reality, the first daughters of aged wildtype mothers also divide slower, which will contribute to differences in colony size, so aged fitness defects are a sum of the slow division times of aged mothers and their first daughters,

For viability assays, log phase cells were inoculated in YPD at 1×10^4^ cells/ml and 60 µl spread on a YPD plate. 1 µM β-estradiol was then added to the cultures, and after 40-48 h 1.5 ml of cells were harvested, washed with 100 µl of water and 40 µl spread on a YPD plate. Colonies were counted after 2 d (young) or 3 d (aged). We note that conclusions from these assays are limited and rely on selecting time points at which replicative ages are matched, this compromise is necessary as culture system has a huge effect on lifespan, with cells in classical microdissection-based lifespan assays living far longer than they do in liquid (47).

### RNA extraction

Cells were re-suspended in 50 µl Lysis/Binding Buffer (from mirVANA kit, Life Technologies AM1560), and 50 µl 0.5 µm zirconium beads (Thistle Scientific 11079105Z) added. Cells were lysed with 5 cycles of 30 s 6500 ms^-1^ / 30 s on ice in Next Advance Bullet Blender Storm 24 (ThermoFisher) in cold room, then 250 µl Lysis/Binding buffer was added followed by 15 µl miRNA Homogenate Additive and cells were briefly vortexed before incubating for 10 minutes on ice. 300 µl acid phenol : chloroform was added, vortexed and centrifuged 5 min at 13,000 g at room temperature before extraction of the upper phase. 400 µl room temperature ethanol and 2 µl glycogen (Sigma G1767) were added and mixture incubated for 1 hour at -30 °C before centrifugation for 15 minutes at 20,000 g, 4 °C. The pellet was washed with cold 70% ethanol and re-suspended in 10 µl water. 1 µl RNA was glyoxylated and analysed on a BPTE mini-gel, and RNA was quantified using a Qubit RNA HS Assay Kit.

150 ng RNA was used to prepare libraries using the NEBNext Ultra II Directional mRNA-seq kit with poly(A)+ purification module (NEB E7760, E7490) as described with modifications: Reagent volumes for elution from poly(T) beads, reverse transcription, second strand synthesis, tailing and adaptor ligation were reduced by 50%; libraries were amplified for 13 cycles using 2 µl each primer per 50 µl reaction before two rounds of AMPure bead purification at 0.9x and elution in 11 µl 0.1x TE prior to quality control using a Bioanalyzer HS DNA ChIP (Agilent) and quantification using a KAPA Library Quantification Kit (Roche).

### DNA extraction and Southern blot analysis

Cell pellets were resuspended in 50 μl 0.34 U/ml lyticase (Sigma L4025) in 1.2 M sorbitol, 50 mM EDTA, 10 mM DTT and incubated at 37°C for 45 min. After centrifugation at 1,000 g for 5 min, cells were gently resuspended in 80 μl of 0.3% SDS, 50 mM EDTA, 250 μg/ml Proteinase K (Roche 3115801) and incubated at 65°C for 30 min. 32 μl 5 M KOAc was added after cooling to room temperature, samples were mixed by flicking, and then chilled on ice for 30 min. After 10 min centrifugation at 20,000 g, the supernatant was extracted into a new tube using a cut tip, 125 μl phenol:chloroform (pH 8) was added and samples were mixed on a wheel for 30 min. Samples were centrifuged for 5 min at 20,000 g, the upper phase was extracted using cut tips, and precipitated with 250 μl ethanol. Pellets were washed with 70% ethanol, air-dried and left overnight at 4°C to dissolve in 20 μl TE. After gentle mixing, 10 μl of each sample was digested with 20 U XhoI (NEB) for 3 to 6 h in 20 μl 1× CutSmart buffer (NEB), 0.2 μl was quantified using PicoGreen DNA (Life Technologies), and equivalent amounts of DNA separated on 25 cm 1% 1× TBE gels overnight at 120 V. Gels were washed in 0.25 N HCl for 15 min, 0.5 N NaOH for 45 min, and twice in 1.5 M NaCl, 0.5 M Tris (pH 7.5) for 20 min before being transferred to 20 × 20 cm HyBond N+ membrane in 6× SSC. Membranes were probed using a biotin-labelled RNA probe to the NTS1 rDNA intergenic spacer region in 10 ml UltraHyb (Life Technologies) at 42°C and washed with 0.1× SSC 0.1% SDS at 42°C, using standard methods.

### Sequencing and bioinformatics

Libraries were sequenced by the Babraham Institute Sequencing Facility using a NextSeq 500 instrument in 75 bp single end mode. After adapter and quality trimming using Trim Galore (v0.6.6), RNA-seq data was mapped to yeast genome R64-1-1 using HISAT2 v2.1.0 90 by the Babraham Institute Bioinformatics Facility. Mapped data was imported into SeqMonk v1.47.0 (https://www.bioinformatics.babraham.ac.uk/projects/seqmonk/) and quantified for log_2_ total reads mapping to the antisense strand of annotated open reading frames (opposite strand specific libraries), excluding the mtDNA and the rDNA locus. Read counts were adjusted by Size Factor normalisation for the full set of quantified probes (91). GO analysis of individual clusters performed using GOrilla (http://cbl-gorilla.cs.technion.ac.il/) (92). Quoted p-values for GO analysis are FDR-corrected according to the Benjamini and Hochberg method (q-values from the GOrilla output), for brevity only the order of magnitude rather than the full q-value is given (93).

### Flow cytometry

Cell pellets were re-suspended in 240 µl PBS and 9 μl 10% triton X-100 containing 0.3 µl of 1 mg/ml Streptavidin conjugated with Alexa Fluor 647 (Life technologies) and 0.6 μl of 1 mg/ml Wheat Germ Agglutinin (WGA) conjugated with CF405S (Biotium). Cells were stained for 10 min at RT on a rotating mixer while covered with aluminium foil, washed once with 300 μl PBS containing 0.01% Triton X-100, re-suspended in 200 μl PBS and immediately subject to flow cytometry analysis on a Cytek Aurora. Cells for single colour controls were prepared in the same manner as the fully stained samples. On the Aurora, time-matched unmixing reference files were required for different ages as cell size and autofluorescence varied with age. Therefore, numerical differences in fluorescence measurements between different ages are not absolutely quantitative. Cell populations were gated on FSH-H and SSC-H to remove debris, on FSC-A and FSC-H to select for single cells, and streptavidin positive (AF647) cells were selected, all in a hierarchical manner and 10,000 events acquired. Low senescence aged populations were defined using quadrants of low GFP and high WGA, with quadrants being defined on the wild-type and applied uniformly across samples within an experiment. As is visible in the representative flow plots, quadrants are almost invariant across most datasets, with differences arising over time through changes in machine performance.

A subset of samples was acquired on an Amnis ImageStream X Mk II with the following laser power settings: 405 = 25 mW, 488 = 180 mW, 561 = 185 mW, 642 = 5 mW, SSC = 0.3 mW. On the ImageStream, cell populations were gated for; (1) single cells based on area and aspect ratio (>0.8), (2) in-focus cells were gated based on a gradient RMS value (>50), and (3) streptavidin positive (AF647) cells, all in a hierarchical manner and 1,000 events acquired. Before data analysis, compensation was applied according to single-colour controls and auto-generated compensation matrix, see (18) for additional parameters.

As absolute measured fluorescence intensities varied somewhat over time and between machines, median normalisation has been applied across all samples of a given age in certain datasets to allow comparison of Tom70-GFP and WGA intensities between datasets. This is a single transformation applied equally to all samples such that differences between samples are maintained, and is not applied to the quadrant analysis described above.

### Oxygen Consumption measurements

Seahorse XF96 cell culture microplates (Agilent) were coated the day before the assay with 100 µL of 1:1 diluted poly-D-lysine (50 µg/mL; Sigma P6407) in ultrapure water for 30 min at room temperature (RT). Excess was removed, plates were air-dried, and stored at 4°C until use. The Seahorse cartridge was hydrated overnight in XF Calibrant at 30°C. Yeast strains for respiratory analyses were grown in Yeast Seahorse Medium (YSM; 0.167% yeast nitrogen base, 0.5% ammonium sulfate, 2% dextrose) with shaking at 30°C. Log-phase cells were cultured overnight to exponential phase and collected on assay day. For aged samples, cells were grown in YPD for 48 h, then live-purified as described above. On the day of the assay, cells were pelleted (3000 rpm, 3 min) and resuspended in YSM. Plates were pre-warmed to 30°C for ≥1 h before seeding ∼5,000 cells per well in 200 μL YSM. Plates were centrifuged (500 rpm, 3 min) to promote adhesion and incubated for 30 min at 30°C for acclimation. OCR was measured using an Agilent Seahorse XF24 Extracellular Flux Analyzer according to the manufacturer’s instructions (30°C). Basal respiration was assessed without drug injections, and average absolute OCR values were recorded. Three sequential absolute OCR measurements were collected in 3 min cycles that include a 1.5 min mix step between measurements.

### Statistical analysis

All statistical analysis was performed in GraphPad Prism (v10.4.1), tests and n values are given in Figure legends.

### Data availability

RNA-seq data is deposited at the Gene Expression Omnibus (GEO), Accession GSE308402.

## Acknowledgements

We would like to thank Sophie Trefely and Mark Cheong for experimental help and insightful discussions. We also thank Sophie Trefely for critical reading of the manuscript. We would like to thank Rachael Walker, Sam Thompson, and Christopher Hall of the Babraham Institute Flow Facility (Flowcyt04) for extensive experimental advice and troubleshooting. RNA-seq library sequencing and processing were performed by Babraham Institute Genomics (Geno06) and Babraham Institute Bioinformatics (Bioinf01), which receive financial support from the Institute Core Capability Grant (BBSRC CCG).

---

**Supplementary Figure S1:**
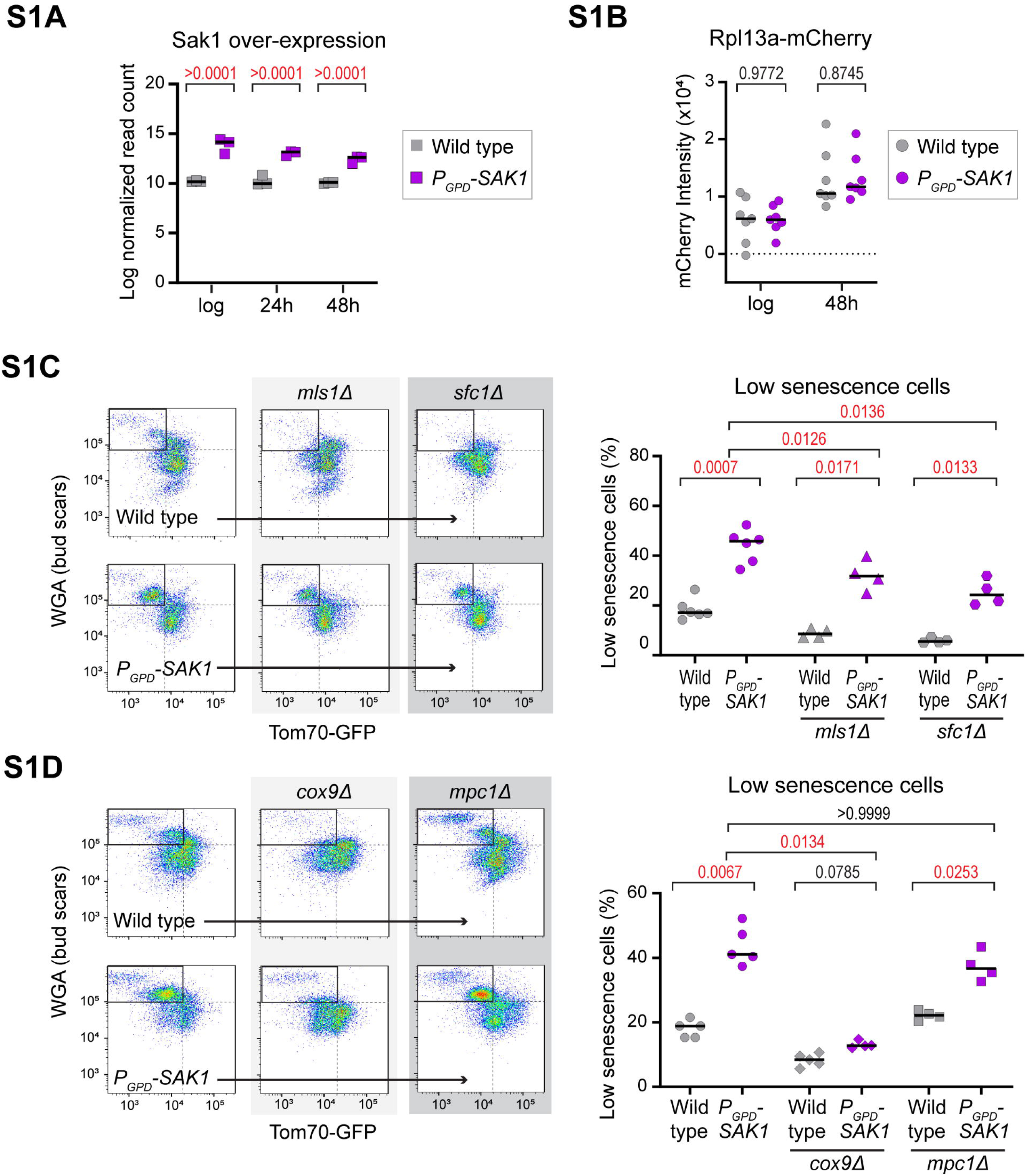
Increased AMPK activity suppresses senescence in a subset of cells. **A:** mRNA expression level of constitutive over-expression of the upstream kinase Sak1 using a P_GPD_ promoter fused to *SAK1* at log phase, 24 h and 48 h. p-values calculated by 1 way ANOVA, n=3 **B:** Impact of *SAK1* overexpression on Rpl13a-mCherry in young and aged cells. p-values calculated by two way ANOVA, n=7 **C:** Representative flow plots and quantification of the low senescence population at 48 h in wild type and *P_GPD_-SAK1* mutants lacking the glyoxylate cycle (*mls1*Δ) or succinate import (*sfc1*Δ). p-values calculated by one way ANOVA, n=4-6 **D:** Representative flow plots and quantification of wild type and *P_GPD_-SAK1* low senescence populations at 48 h in mitochondrial mutants lacking a functional electron transport chain (*cox9*Δ) or pyruvate import (*mpc1*Δ). p-values calculated by one way ANOVA, n=4-5

**Supplementary Figure S2:**
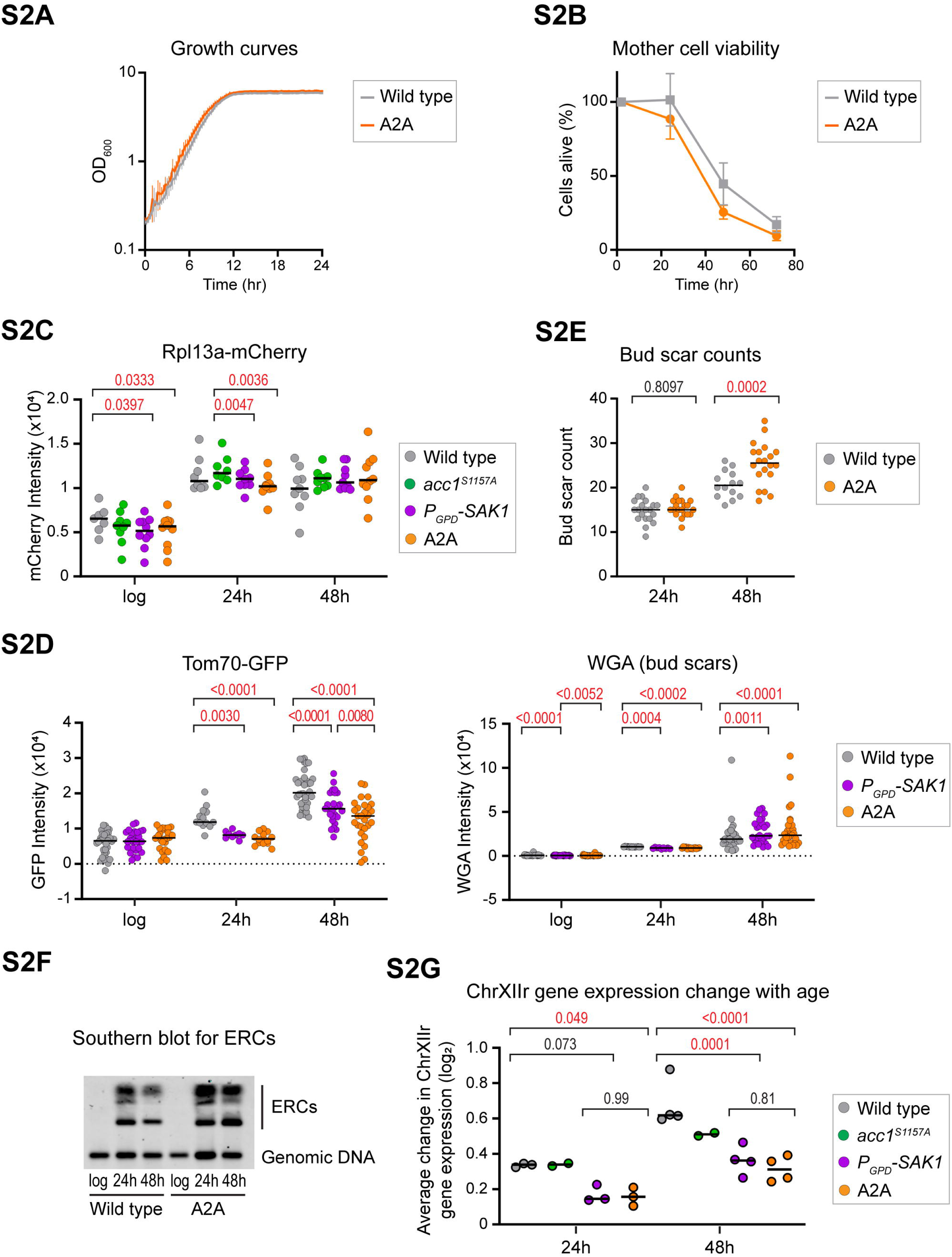
Combining AMPK activity and fatty acid synthesis to avoid senescence. **A:** Growth curve of wild type and A2A cells in YPD media starting from log-phase pre-cultures. **B:** Lifespan measured in YPD media for wild type and A2A cells based on %age of viable cells remaining at each time point; *n* = 5. **C:** Population medians for Rpl13a-mCherry in wild type, *acc1^S1157A^*, *P_GPD_-SAK1* and A2A at log phase, 24 h and 48 h. p values calculated by two way ANOVA, n=7-10. **D:** Population medians for Tom70-GFP and WGA (bud scars) in wild type, *P_GPD_-SAK1,* and A2A at log phase, 24 h and 48 h made from a compilation of experiments performed during this study. p values calculated by two way ANOVA, n=27-38. **E:** Manual bud scar counts of log phase and 48 h-aged wild type and A2A cells. p values calculated by one way ANOVA, n=14-20 **F**: Southern blot analysis of ERCs compared to genomic rDNA in wild type and A2A cells at log, 24 h and 48 h. **G**: Analysis of median change in log2 mRNA abundance from log phase to given time points for all genes on ChrXIIr. Analysis by 2-way ANOVA for aged time-points only since log values are the reference point n = 2-4 biological replicates.

**Supplementary Figure S3:**
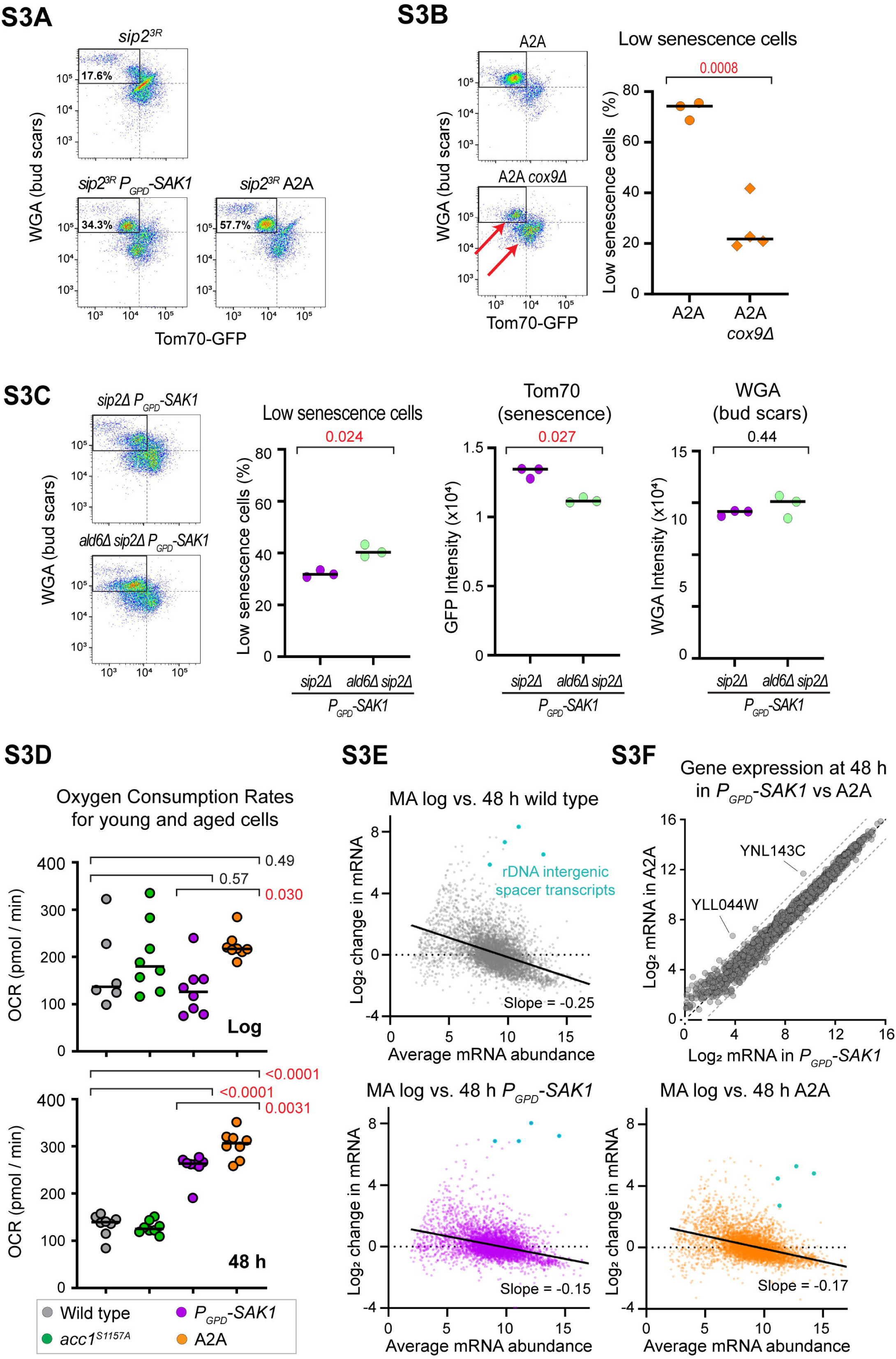
Genetic evidence that cytosolic Acetyl-CoA promotes senescence. **A:** Representative flow plots of the low senescence population at 48 h in *sip2^3R^*, *sip2^3R^ P_GPD_-SAK1* and *sip2^3R^*A2A. **B:** Representative flow plots and quantification of low senescence population at 48 h in A2A mutants lacking the electron transport chain (*cox9*Δ), p-value calculated by t-test, n=3-4. **C:** Representative flow plots and quantification of low senescence population in wild type and *sip2*Δ *P_GPD_-SAK1 ald6*Δ at 48 h. Population medians are also shown for Tom70-GFP and WGA in wild type and *ald6*Δ at 48 h. p values calculated by t-test, n=5-6 **D:** Basal oxygen consumption rates for log and aged cells. p values calculated by one way ANOVA, n = 6-8. Note that due to unavoidable differences in cell number, the values for log and 48 h samples are not comparable. **E:** MA plots comparing change in normalised mRNA abundance between log and 48 h to average mRNA abundance between log and 48 h. Data represents an average of 3-4 biological replicate RNA-seq experiments. Line of best fit was calculated by linear regression, with the slope parameter quoted as a measure of gene expression dysregulation (18). Transcripts from the rDNA intergenic spacer are highlighted in blue. **F:** Comparison of gene expression profiles of *P_GPD_-SAK1* and A2A cells at 48 h ageing by mRNA-seq. Data is an average of 4 biological replicates, there are no significantly different genes (p<0.05 by DEseq2). 3-fold differences are indicated by dotted lines, only uncharacterised genes pass this threshold and are not significant.

**Supplementary Figure S4:**
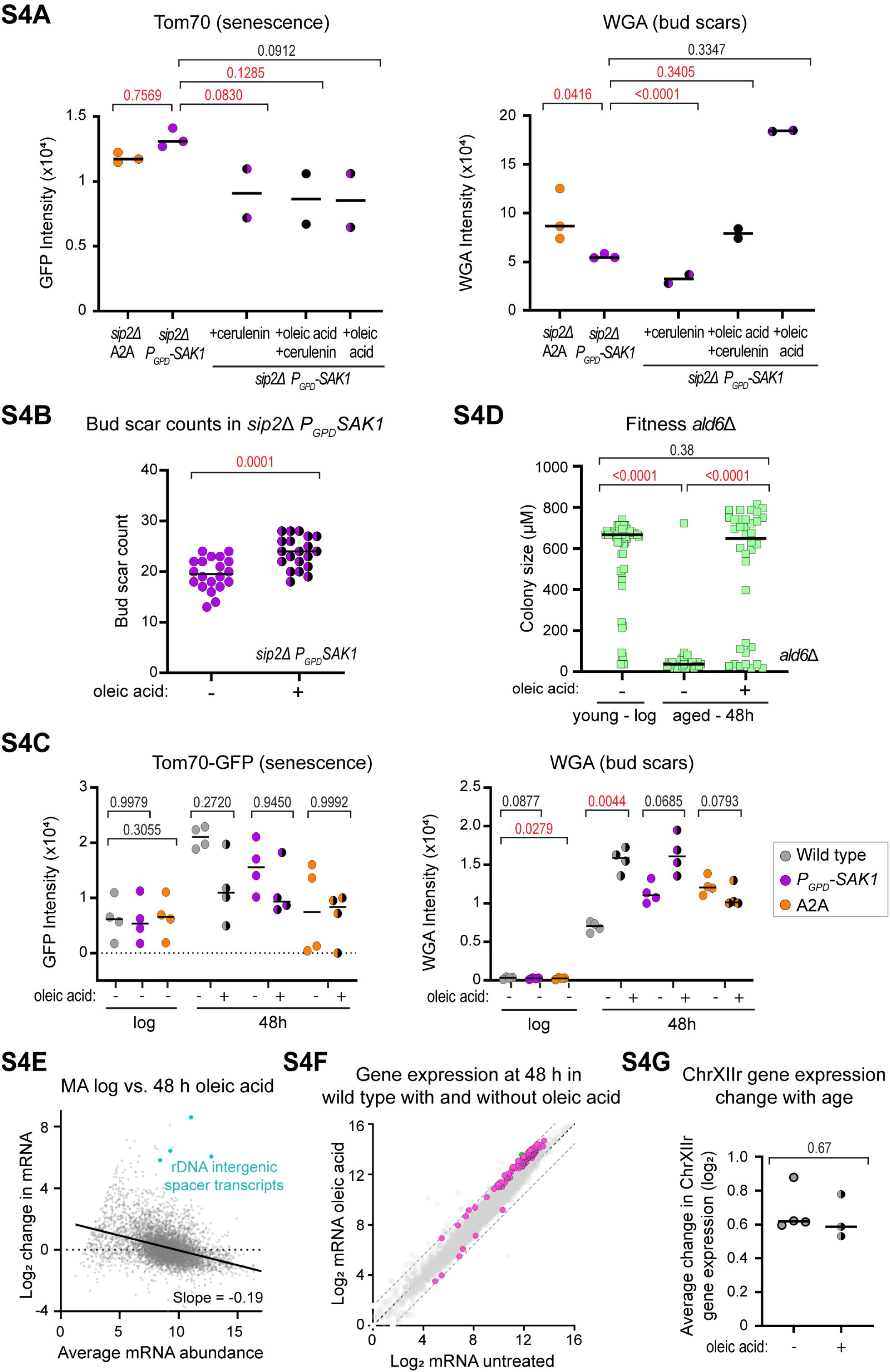
Fatty acids in senescence. **A:** Population medians for Tom70-GFP and WGA at 48 h in *sip2*Δ *P_GPD_-SAK1 acc1^S1157A^* and *sip2*Δ *P_GPD_-SAK1* supplemented with cerulenin and/or 0.04% oleic acid + 0.04% Tween 80. p-values calculated by one-way ANOVA, n=2-3 **B:** Manual bud scar counts of 48 h-aged *sip2*Δ *P_GPD_-SAK1* cells with and without 0.04% oleic acid. p values calculated by one way ANOVA, n=20 **C:** Population medians for Tom70-GFP and WGA at 48 h in wild type, *P_GPD_-SAK1* and A2A with and without 0.04% oleic acid + 0.04% Tween 80. p-values calculated by one-way ANOVA, n=4 **D:** Size of colonies formed in 24 h on a YPD plate by log phase and 48 h-aged *ald6*Δ cells with 0.04% oleic acid + 0.04% Tween 80. Only cells that eventually formed colonies within 3 days were included to ensure all tested cells were viable. p values calculated by one-way ANOVA, n=28-53 **E:** MA plot comparing change in normalised mRNA abundance between log and 48 h to average mRNA abundance between log and 48 h. Data represents an average of 3-4 biological replicate RNA-seq experiments. Line of best fit was calculated by linear regression, with the slope parameter quoted as a measure of gene expression dysregulation (18). Transcripts from the rDNA intergenic spacer are highlighted in blue. **F**: Comparison of gene expression profiles of wild type cells after 48 h ageing with or without 0.04% oleic acid + 0.04% Tween 80 by mRNA-seq. Data is an average of 4 biological replicates (untreated) or 3 biological replicates (oleic acid), significantly different genes (p<0.01 by DEseq2) are shown in purple, genes both substantially (>3x indicated by dotted lines) and significantly different are highlighted in green. **G**: Analysis of median change in log2 mRNA abundance from log phase to 48 h for all genes on ChrXIIr. Analysis by t-test for aged time-points only since log values are the reference point n = 3-4 biological replicates.

## References

1. Medawar, P.B. (1952) An unsolved problem of biology. Published for the college by H. K. Lewis, London,.

2. Kirkwood, T.B. (1977) Evolution of ageing. Nature, 270, 301–304.

3. Williams, G.C. (1957) Pleiotropy, natural selection and the evolution of senescence. Evolution, 11, 398–411.

4. Bylino, O.V., Ogienko, A.A., Batin, M.A., Georgiev, P.G. and Omelina, E.S. (2024) Genetic, Environmental, and Stochastic Components of Lifespan Variability: The Drosophila Paradigm. Int J Mol Sci, 25.

5. Brooks-Wilson, A.R. (2013) Genetics of healthy aging and longevity. Hum Genet, 132, 1323–1338.

6. Li, Y., Jiang, Y., Paxman, J., O’Laughlin, R., Klepin, S., Zhu, Y., Pillus, L., Tsimring, L.S., Hasty, J. and Hao, N. (2020) A programmable fate decision landscape underlies single-cell aging in yeast. Science, 369, 325–329.

7. Speakman, J.R. and Mitchell, S.E. (2011) Caloric restriction. Mol Aspects Med, 32, 159–221.

8. Osborne, T.B., Mendel, L.B. and Ferry, E.L. (1917) The Effect of Retardation of Growth Upon the Breeding Period and Duration of Life of Rats. Science, 45, 294–295.

9. Colman, R.J., Anderson, R.M., Johnson, S.C., Kastman, E.K., Kosmatka, K.J., Beasley, T.M., Allison, D.B., Cruzen, C., Simmons, H.A., Kemnitz, J.W. et al. (2009) Caloric restriction delays disease onset and mortality in rhesus monkeys. Science, 325, 201–204.

10. Mattison, J.A., Roth, G.S., Beasley, T.M., Tilmont, E.M., Handy, A.M., Herbert, R.L., Longo, D.L., Allison, D.B., Young, J.E., Bryant, M. et al. (2012) Impact of caloric restriction on health and survival in rhesus monkeys from the NIA study. Nature, 489, 318–321.

11. Mannick, J.B. and Lamming, D.W. (2023) Targeting the biology of aging with mTOR inhibitors. Nat Aging, 3, 642–660.

12. Mortimer, R.K. and Johnston, J.R. (1959) Life span of individual yeast cells. Nature, 183, 1751–1752.

13. Egilmez, N.K. and Jazwinski, S.M. (1989) Evidence for the involvement of a cytoplasmic factor in the aging of the yeast Saccharomyces cerevisiae. Journal of bacteriology, 171, 37–42.

14. Fehrmann, S., Paoletti, C., Goulev, Y., Ungureanu, A., Aguilaniu, H. and Charvin, G. (2013) Aging yeast cells undergo a sharp entry into senescence unrelated to the loss of mitochondrial membrane potential. Cell reports, 5, 1589–1599.

15. Morlot, S., Song, J., Leger-Silvestre, I., Matifas, A., Gadal, O. and Charvin, G. (2019) Excessive rDNA Transcription Drives the Disruption in Nuclear Homeostasis during Entry into Senescence in Budding Yeast. Cell reports, 28, 408–422 e404.

16. Jin, M., Li, Y., O’Laughlin, R., Bittihn, P., Pillus, L., Tsimring, L.S., Hasty, J. and Hao, N. (2019) Divergent Aging of Isogenic Yeast Cells Revealed through Single-Cell Phenotypic Dynamics. Cell Syst, 8, 242–253 e243.

17. Wang, J., Sang, Y., Jin, S., Wang, X., Azad, G.K., McCormick, M.A., Kennedy, B.K., Li, Q., Wang, J., Zhang, X. et al. (2022) Single-cell RNA-seq reveals early heterogeneity during aging in yeast. Aging cell, 21, e13712.

18. Horkai, D., Hadj-Moussa, H., Whale, A.J. and Houseley, J. (2023) Dietary change without caloric restriction maintains a youthful profile in ageing yeast. PLoS Biol, 21, e3002245.

19. Herzig, S. and Shaw, R.J. (2018) AMPK: guardian of metabolism and mitochondrial homeostasis. Nat Rev Mol Cell Biol, 19, 121–135.

20. Garcia, D. and Shaw, R.J. (2017) AMPK: Mechanisms of Cellular Energy Sensing and Restoration of Metabolic Balance. Mol Cell, 66, 789–800.

21. Salminen, A. and Kaarniranta, K. (2012) AMP-activated protein kinase (AMPK) controls the aging process via an integrated signaling network. Ageing Res Rev, 11, 230–241.

22. Burkewitz, K., Zhang, Y. and Mair, W.B. (2014) AMPK at the nexus of energetics and aging. Cell metabolism, 20, 10–25.

23. Wierman, M.B., Maqani, N., Strickler, E., Li, M. and Smith, J.S. (2017) Caloric Restriction Extends Yeast Chronological Life Span by Optimizing the Snf1 (AMPK) Signaling Pathway. Mol Cell Biol, 37.

24. Stenesen, D., Suh, J.M., Seo, J., Yu, K., Lee, K.S., Kim, J.S., Min, K.J. and Graff, J.M. (2013) Adenosine nucleotide biosynthesis and AMPK regulate adult life span and mediate the longevity benefit of caloric restriction in flies. Cell metabolism, 17, 101–112.

25. Greer, E.L., Dowlatshahi, D., Banko, M.R., Villen, J., Hoang, K., Blanchard, D., Gygi, S.P. and Brunet, A. (2007) An AMPK-FOXO pathway mediates longevity induced by a novel method of dietary restriction in C. elegans. Curr Biol, 17, 1646–1656.

26. Martin-Montalvo, A., Mercken, E.M., Mitchell, S.J., Palacios, H.H., Mote, P.L., Scheibye-Knudsen, M., Gomes, A.P., Ward, T.M., Minor, R.K., Blouin, M.J. et al. (2013) Metformin improves healthspan and lifespan in mice. Nature communications, 4, 2192.

27. Mair, W., Morantte, I., Rodrigues, A.P., Manning, G., Montminy, M., Shaw, R.J. and Dillin, A. (2011) Lifespan extension induced by AMPK and calcineurin is mediated by CRTC-1 and CREB. Nature, 470, 404–408.

28. Ulgherait, M., Rana, A., Rera, M., Graniel, J. and Walker, D.W. (2014) AMPK modulates tissue and organismal aging in a non-cell-autonomous manner. Cell reports, 8, 1767–1780.

29. Apfeld, J., O’Connor, G., McDonagh, T., DiStefano, P.S. and Curtis, R. (2004) The AMP-activated protein kinase AAK-2 links energy levels and insulin-like signals to lifespan in C. elegans. Genes Dev, 18, 3004–3009.

30. Yang, S., Long, L.H., Li, D., Zhang, J.K., Jin, S., Wang, F. and Chen, J.G. (2015) beta-Guanidinopropionic acid extends the lifespan of Drosophila melanogaster via an AMP-activated protein kinase-dependent increase in autophagy. Aging cell, 14, 1024–1033.

31. Mohammed, I., Hollenberg, M.D., Ding, H. and Triggle, C.R. (2021) A Critical Review of the Evidence That Metformin Is a Putative Anti-Aging Drug That Enhances Healthspan and Extends Lifespan. Front Endocrinol (Lausanne), 12, 718942.

32. Slack, C., Foley, A. and Partridge, L. (2012) Activation of AMPK by the putative dietary restriction mimetic metformin is insufficient to extend lifespan in Drosophila. PLoS One, 7, e47699.

33. Jiang, J.C., Jaruga, E., Repnevskaya, M.V. and Jazwinski, S.M. (2000) An intervention resembling caloric restriction prolongs life span and retards aging in yeast. FASEB journal : official publication of the Federation of American Societies for Experimental Biology, 14, 2135–2137.

34. Anderson, R.M., Bitterman, K.J., Wood, J.G., Medvedik, O. and Sinclair, D.A. (2003) Nicotinamide and PNC1 govern lifespan extension by calorie restriction in Saccharomyces cerevisiae. Nature, 423, 181–185.

35. Kaeberlein, M., Kirkland, K.T., Fields, S. and Kennedy, B.K. (2004) Sir2-independent life span extension by calorie restriction in yeast. PLoS Biol, 2, E296.

36. Maqani, N., Fine, R.D., Shahid, M., Li, M., Enriquez-Hesles, E. and Smith, J.S. (2018) Spontaneous mutations in CYC8 and MIG1 suppress the short chronological lifespan of budding yeast lacking SNF1/AMPK. Microb Cell, 5, 233–248.

37. Ashrafi, K., Lin, S.S., Manchester, J.K. and Gordon, J.I. (2000) Sip2p and its partner snf1p kinase affect aging in S. cerevisiae. Genes Dev, 14, 1872–1885.

38. Simpson-Lavy, K.J. and Kupiec, M. (2022) Regulation of yeast Snf1 (AMPK) by a polyhistidine containing pH sensing module. iScience, 25, 105083.

39. Rubenstein, E.M., McCartney, R.R., Zhang, C., Shokat, K.M., Shirra, M.K., Arndt, K.M. and Schmidt, M.C. (2008) Access denied: Snf1 activation loop phosphorylation is controlled by availability of the phosphorylated threonine 210 to the PP1 phosphatase. J Biol Chem, 283, 222–230.

40. Hong, S.P., Leiper, F.C., Woods, A., Carling, D. and Carlson, M. (2003) Activation of yeast Snf1 and mammalian AMP-activated protein kinase by upstream kinases. PNAS, 100, 8839–8843.

41. Pan, D.A. and Hardie, D.G. (2002) A homologue of AMP-activated protein kinase in Drosophila melanogaster is sensitive to AMP and is activated by ATP depletion. The Biochemical journal, 367, 179–186.

42. Munday, M.R., Campbell, D.G., Carling, D. and Hardie, D.G. (1988) Identification by amino acid sequencing of three major regulatory phosphorylation sites on rat acetyl-CoA carboxylase. Eur J Biochem, 175, 331–338.

43. Woods, A., Munday, M.R., Scott, J., Yang, X., Carlson, M. and Carling, D. (1994) Yeast SNF1 is functionally related to mammalian AMP-activated protein kinase and regulates acetyl-CoA carboxylase in vivo. J Biol Chem, 269, 19509–19515.

44. Davies, S.P., Sim, A.T. and Hardie, D.G. (1990) Location and function of three sites phosphorylated on rat acetyl-CoA carboxylase by the AMP-activated protein kinase. Eur J Biochem, 187, 183–190.

45. Neigeborn, L. and Carlson, M. (1984) Genes affecting the regulation of SUC2 gene expression by glucose repression in Saccharomyces cerevisiae. Genetics, 108, 845–858.

46. Carlson, M., Osmond, B.C. and Botstein, D. (1981) Mutants of yeast defective in sucrose utilization. Genetics, 98, 25–40.

47. Lindstrom, D.L. and Gottschling, D.E. (2009) The mother enrichment program: a genetic system for facile replicative life span analysis in Saccharomyces cerevisiae. Genetics, 183, 413–422.

48. Lindstrom, D.L., Leverich, C.K., Henderson, K.A. and Gottschling, D.E. (2011) Replicative age induces mitotic recombination in the ribosomal RNA gene cluster of Saccharomyces cerevisiae. PLoS Genet, 7, e1002015.

49. Allen, C., Buttner, S., Aragon, A.D., Thomas, J.A., Meirelles, O., Jaetao, J.E., Benn, D., Ruby, S.W., Veenhuis, M., Madeo, F. et al. (2006) Isolation of quiescent and nonquiescent cells from yeast stationary-phase cultures. The Journal of cell biology, 174, 89–100.

50. Zylstra, A., Hadj-Moussa, H., Horkai, D., Whale, A.J., Piguet, B. and Houseley, J. (2023) Senescence in yeast is associated with amplified linear fragments of chromosome XII rather than ribosomal DNA circle accumulation. PLoS Biol, 21, e3002250.

51. Patterson, M.N. and Maxwell, P.H. (2014) Combining magnetic sorting of mother cells and fluctuation tests to analyze genome instability during mitotic cell aging in Saccharomyces cerevisiae. *Journal of visualized experiments : JoVE*, e51850.

52. Janssens, G.E., Meinema, A.C., Gonzalez, J., Wolters, J.C., Schmidt, A., Guryev, V., Bischoff, R., Wit, E.C., Veenhoff, L.M. and Heinemann, M. (2015) Protein biogenesis machinery is a driver of replicative aging in yeast. eLife, 4, e08527.

53. Chen, C. and Contreras, R. (2007) Identifying genes that extend life span using a high-throughput screening system. Methods Mol Biol, 371, 237–248.

54. Janssens, G.E. and Veenhoff, L.M. (2016) The Natural Variation in Lifespans of Single Yeast Cells Is Related to Variation in Cell Size, Ribosomal Protein, and Division Time. PLoS One, 11, e0167394.

55. Raab, A.M., Hlavacek, V., Bolotina, N. and Lang, C. (2011) Shifting the fermentative/oxidative balance in Saccharomyces cerevisiae by transcriptional deregulation of Snf1 via overexpression of the upstream activating kinase Sak1p. Applied and environmental microbiology, 77, 1981–1989.

56. Garcia-Salcedo, R., Lubitz, T., Beltran, G., Elbing, K., Tian, Y., Frey, S., Wolkenhauer, O., Krantz, M., Klipp, E. and Hohmann, S. (2014) Glucose de-repression by yeast AMP-activated protein kinase SNF1 is controlled via at least two independent steps. FEBS J, 281, 1901–1917.

57. Schuller, H.J. (2003) Transcriptional control of nonfermentative metabolism in the yeast Saccharomyces cerevisiae. Curr Genet, 43, 139–160.

58. Gancedo, J.M. (1998) Yeast carbon catabolite repression. Microbiology and molecular biology reviews : MMBR, 62, 334–361.

59. Bojunga, N., Kotter, P. and Entian, K.D. (1998) The succinate/fumarate transporter Acr1p of Saccharomyces cerevisiae is part of the gluconeogenic pathway and its expression is regulated by Cat8p. Molecular & general genetics : MGG, 260, 453–461.

60. Zhou, Z., Liu, Y., Feng, Y., Klepin, S., Tsimring, L.S., Pillus, L., Hasty, J. and Hao, N. (2023) Engineering longevity-design of a synthetic gene oscillator to slow cellular aging. Science, 380, 376–381.

61. Shi, S., Chen, Y., Siewers, V. and Nielsen, J. (2014) Improving production of malonyl coenzyme A-derived metabolites by abolishing Snf1-dependent regulation of Acc1. mBio, 5, e01130–01114.

62. Choi, J.W. and Da Silva, N.A. (2014) Improving polyketide and fatty acid synthesis by engineering of the yeast acetyl-CoA carboxylase. Journal of biotechnology, 187, 56–59.

63. Sinclair, D.A. and Guarente, L. (1997) Extrachromosomal rDNA circles--a cause of aging in yeast. Cell, 91, 1033–1042.

64. Sinclair, D.A., Mills, K. and Guarente, L. (1997) Accelerated aging and nucleolar fragmentation in yeast sgs1 mutants. Science, 277, 1313–1316.

65. Lu, J.Y., Lin, Y.Y., Sheu, J.C., Wu, J.T., Lee, F.J., Chen, Y., Lin, M.I., Chiang, F.T., Tai, T.Y., Berger, S.L. et al. (2011) Acetylation of yeast AMPK controls intrinsic aging independently of caloric restriction. Cell, 146, 969–979.

66. Gross, A.S., Zimmermann, A., Pendl, T., Schroeder, S., Schoenlechner, H., Knittelfelder, O., Lamplmayr, L., Santiso, A., Aufschnaiter, A., Waltenstorfer, D. et al. (2019) Acetyl-CoA carboxylase 1-dependent lipogenesis promotes autophagy downstream of AMPK. J Biol Chem, 294, 12020–12039.

67. Shiba, Y., Paradise, E.M., Kirby, J., Ro, D.K. and Keasling, J.D. (2007) Engineering of the pyruvate dehydrogenase bypass in Saccharomyces cerevisiae for high-level production of isoprenoids. Metabolic engineering, 9, 160–168.

68. Meaden, P.G., Dickinson, F.M., Mifsud, A., Tessier, W., Westwater, J., Bussey, H. and Midgley, M. (1997) The ALD6 gene of Saccharomyces cerevisiae encodes a cytosolic, Mg(2+)-activated acetaldehyde dehydrogenase. Yeast, 13, 1319–1327.

69. Hu, Z., Chen, K., Xia, Z., Chavez, M., Pal, S., Seol, J.H., Chen, C.C., Li, W. and Tyler, J.K. (2014) Nucleosome loss leads to global transcriptional up-regulation and genomic instability during yeast aging. Genes Dev, 28, 396–408.

70. Lesur, I. and Campbell, J.L. (2004) The transcriptome of prematurely aging yeast cells is similar to that of telomerase-deficient cells. Molecular biology of the cell, 15, 1297–1312.

71. Yiu, G., McCord, A., Wise, A., Jindal, R., Hardee, J., Kuo, A., Shimogawa, M.Y., Cahoon, L., Wu, M., Kloke, J. et al. (2008) Pathways change in expression during replicative aging in Saccharomyces cerevisiae. The journals of gerontology. Series A, Biological sciences and medical sciences, 63, 21–34.

72. Nicastro, R., Brohee, L., Alba, J., Nuchel, J., Figlia, G., Kipschull, S., Gollwitzer, P., Romero-Pozuelo, J., Fernandes, S.A., Lamprakis, A. et al. (2023) Malonyl-CoA is a conserved endogenous ATP-competitive mTORC1 inhibitor. Nat Cell Biol, 25, 1303–1318.

73. Kawaguchi, A., Tomoda, H., Nozoe, S., Omura, S. and Okuda, S. (1982) Mechanism of action of cerulenin on fatty acid synthetase. Effect of cerulenin on iodoacetamide-induced malonyl-CoA decarboxylase activity. Journal of biochemistry, 92, 7–12.

74. Bagamery, L.E., Justman, Q.A., Garner, E.C. and Murray, A.W. (2020) A Putative Bet-Hedging Strategy Buffers Budding Yeast against Environmental Instability. Curr Biol, 30, 4563–4578 e4564.

75. Levy, S.F., Ziv, N. and Siegal, M.L. (2012) Bet hedging in yeast by heterogeneous, age-correlated expression of a stress protectant. PLoS Biol, 10, e1001325.

76. Frenk, S., Pizza, G., Walker, R.V. and Houseley, J. (2017) Aging yeast gain a competitive advantage on non-optimal carbon sources. Aging cell, 16, 602–604.

77. Moreno, D.F., Jenkins, K., Morlot, S., Charvin, G., Csikasz-Nagy, A. and Aldea, M. (2019) Proteostasis collapse, a hallmark of aging, hinders the chaperone-Start network and arrests cells in G1. eLife, 8.

78. Neurohr, G.E., Terry, R.L., Sandikci, A., Zou, K., Li, H. and Amon, A. (2018) Deregulation of the G1/S-phase transition is the proximal cause of mortality in old yeast mother cells. Genes Dev, 32, 1075–1084.

79. Huberts, D.H., Gonzalez, J., Lee, S.S., Litsios, A., Hubmann, G., Wit, E.C. and Heinemann, M. (2014) Calorie restriction does not elicit a robust extension of replicative lifespan in Saccharomyces cerevisiae. PNAS, 111, 11727–11731.

80. Jo, M.C., Liu, W., Gu, L., Dang, W. and Qin, L. (2015) High-throughput analysis of yeast replicative aging using a microfluidic system. PNAS.

81. Kwon, Y.Y., Lee, S.K. and Lee, C.K. (2017) Caloric Restriction-Induced Extension of Chronological Lifespan Requires Intact Respiration in Budding Yeast. Mol Cells, 40, 307–313.

82. Lin, S.S., Manchester, J.K. and Gordon, J.I. (2003) Sip2, an N-myristoylated beta subunit of Snf1 kinase, regulates aging in Saccharomyces cerevisiae by affecting cellular histone kinase activity, recombination at rDNA loci, and silencing. J Biol Chem, 278, 13390–13397.

83. Lin, S.S., Manchester, J.K. and Gordon, J.I. (2001) Enhanced gluconeogenesis and increased energy storage as hallmarks of aging in Saccharomyces cerevisiae. J Biol Chem, 276, 36000–36007.

84. Fullerton, M.D., Galic, S., Marcinko, K., Sikkema, S., Pulinilkunnil, T., Chen, Z.P., O’Neill, H.M., Ford, R.J., Palanivel, R., O’Brien, M. et al. (2013) Single phosphorylation sites in Acc1 and Acc2 regulate lipid homeostasis and the insulin-sensitizing effects of metformin. Nat Med, 19, 1649–1654.

85. Houseley, J. and Tollervey, D. (2011) Repeat expansion in the budding yeast ribosomal DNA can occur independently of the canonical homologous recombination machinery. Nucleic Acids Res, 39, 8778–8791.

86. Janke, C., Magiera, M.M., Rathfelder, N., Taxis, C., Reber, S., Maekawa, H., Moreno-Borchart, A., Doenges, G., Schwob, E., Schiebel, E. et al. (2004) A versatile toolbox for PCR-based tagging of yeast genes: new fluorescent proteins, more markers and promoter substitution cassettes. Yeast, 21, 947–962.

87. Longtine, M.S., McKenzie, A., 3rd, Demarini, D.J., Shah, N.G., Wach, A., Brachat, A., Philippsen, P. and Pringle, J.R. (1998) Additional modules for versatile and economical PCR-based gene deletion and modification in Saccharomyces cerevisiae. Yeast, 14, 953–961.

88. Storici, F. and Resnick, M.A. (2006) The delitto perfetto approach to in vivo site-directed mutagenesis and chromosome rearrangements with synthetic oligonucleotides in yeast. Methods Enzymol, 409, 329–345.

89. Cruz, C., Della Rosa, M., Krueger, C., Gao, Q., Horkai, D., King, M., Field, L. and Houseley, J. (2018) Tri-methylation of histone H3 lysine 4 facilitates gene expression in ageing cells. eLife, 7, e34081.

90. Kim, D., Langmead, B. and Salzberg, S.L. (2015) HISAT: a fast spliced aligner with low memory requirements. Nat Methods, 12, 357–360.

91. Anders, S. and Huber, W. (2010) Differential expression analysis for sequence count data. Genome Biol, 11, R106.

92. Eden, E., Navon, R., Steinfeld, I., Lipson, D. and Yakhini, Z. (2009) GOrilla: a tool for discovery and visualization of enriched GO terms in ranked gene lists. BMC bioinformatics, 10, 48.

93. Benjamini, Y. and Hochberg, Y. (1995) Controlling the False Discovery Rate - a Practical and Powerful Approach to Multiple Testing. Journal of the Royal Statistical Society Series B-Statistical Methodology, 57, 289–300.

