## Supplementary material for "Decoupling AMPK from fatty acid synthesis allows maintenance of fitness late in life": Table S2

**Table S2: Oligonucleotides used for making strains.**

| oJH1520 | TOM70 DN45 1  (GFP tag) | TAGTTTTTGTCTTCTCCTAAAAGTTTTTAAGTTTATGTTTACTGT GAATTCGAGCTCGTTTAAAC |
| --- | --- | --- |
| oJH1521 | **TOM70 UP45 1**  (GFP tag) | AAGATTCAAGAAACTTTAGCTAAATTACGCGAACAGGGTTTAATG CGG ATC CCC GGG TTA ATT AAC |
| oDH11 | **RPL13A-mCherry UP45**  (for mCherry tag with pAW8) | AAGAGAGCTAGAGAAAAGGCTGAAGCTGAAGCTGAAAAGAAGAAATGCATGCTTATGGTGAGCAA |
| oDH12 | **RPL13A-mCherry DN45**  (for mCherry tag with pAW8) | ATACAAAAATTGTGGATGAAAAATTCTTTGATGAAGTTTTTAGATAAGTTATACTAGTTCGTCGACTGGAT |
| oJH1777 | **ACC1 CORE UP45 1**  (for making S1157A mutation in ACC1) | TTCTCCACCTTTCCAACTGTTAAATCTAAAATGGGTATGAACAGGGCTGT TAGGGATAACAGGGTAAT CCGCGCGTTGGCCGATTCAT |
| oJH1778 | **ACC1 CORE DN45 1**  (for making S1157A mutation in ACC1) | TCTTAACGGAGATGACTGACTGTTTGCAACATATGACAAATCTGAAACAG TTCGTACGCTGCAGGTCGAC |
| oDH31 | **SAK1 over-expression UP45**  (pYM-14 GDP promoter insertion 201bp up from start of SAK1) | TATCCAGTCACAATTTTAAAAGCACTTCGTTAACACGTTTGGTTG CGTACGCTGCAGGTCGAC |
| oDH32 | **SAK1 over-expression DN45**  (pYM-14 GDP promoter insertion 45bp up from start of SAK1 start codon ATG) | AGGTACATTGACCTCTTCGACGTTAACTTTTTTATCACTCCTATC CATCGATGAATTCTCTGTCG |
| oDH113 | **COX9 UP45**  (COX9 deletion with pFA6a-URA plasmid) | AGCAAGATATTTGCAAACTACTAACTACAAGCGACTTACACAGAC CGGATCCCCGGGTTAATTAAG |
| oDH114 | **COX9 DN45**  (COX9 deletion with pFA6a-URA plasmid) | TAGGAATAAGAATATAATGCGAAAAACAATAGTGGTCAGGTTCGG GAATTCGAGCTCGTTTAAAC |
| oMU8 | **MPC1 UP45**  (cloning primers for making MPC1-HIS cassette using pFA6A HisMX6) | TCTACATATATACGTATAGATTTTATTGCACTGTGATCAAAAAGACGGATCCCCGGGTTAATTAAG |
| oMU9 | **MPC1 DN45**  (cloning primers for making MPC1-HIS cassette using pFA6a HisMX6) | ATGCATGAACATATCCATCCTCTTAATCCTTGCTTGTTCTTTTTTGAATTCGAGCTCGTTTAAAC |
| oHH25 | **MLS1 UP45**  (Forward overlap delete MLS1 [(45bp of left flank) + F1 from pFA6a]) | TTTGAACTAAACAAAGTAGTAAAAGCACATAAAAGAATTAAGAAA CGGATCCCCGGGTTAATTAAG |
| oHH26 | **MLS1 DN45**  (Reverse overlap to delete MLS1 [(45bp of right flank) + R1 from pFA6a]) | GGCATGAATATATTTTTATATATGTGTACACTGGGGCAAGGGAGA GAATTCGAGCTCGTTTAAAC |
| oHH33 | **SFC1 UP45**  (Forward overlap delete SFC1 [(45bp of left flank) + F1 from pFA6a]) | TTAGTTTTAGCCGTAAGATAACATAACAAAGAAGAAGAAAGAAAA CGGATCCCCGGGTTAATTAAG |
| oHH34 | **SFC1 DN45**  (Reverse overlap to delete SFC1 [(45bp of right flank) + R1 from pFA6a]) | TTCTATTTTTTCTTTATTTCATTTTGTAGTCCCATTGTTCATCAT GAATTCGAGCTCGTTTAAAC |
| oMU13 | **CAT2 UP45**  (cloning primers for making CAT2-HIS cassette using pFA6A HisMX6) | GTATATATATATATATATCCCTTAAAAACTAAAAAGAAAGCACTCCGGATCCCCGGGTTAATTAAG |
| oMU14 | **CAT2 DN45**  (cloning primers or making CAT2-HIS cassette using pFA6a HisMX6) | TAACAAAAATATTCACAAATTAATTGAAGAGGAAAGGTGAAAAATGAATTCGAGCTCGTTTAAAC |
| oHH96 | **Sip2 UP45**  (overlap UP45 upstream region for deletion of Sip2 with pFA6a URA3) | AAATAATTGTGCTTTTGGAAGGGTACTGAGAGCAGTGAAGTTTCTCGGATCCCCGGGTTAATTAAG |
| oHH97 | **Sip2 DN45**  (overlap DN45 downstream region for deletion of Sip2 with pFA6a URA3) | GCCAAAAATATATACAATACGCCTTTAAAAAGTAGGAATCCCGGAGAATTCGAGCTCGTTTAAAC |
