## Supplementary material for "Decoupling AMPK from fatty acid synthesis allows maintenance of fitness late in life": Table S1

**Table S1: Strains used in this work.**

All strains are diploid derivatives of the MEP system (47). TOM70-GFP and RPL13a-mCherry markers are heterozygous to avoid growth defect.

| DH18 | WT-RPL13a | *ade2::hisG his3 leu2 met15D::ADE2/MET15 lys2/LYS2 ura3DO trp1D63 hoD::SCW11pr-Cre-EBD78-NatMX loxP-UBC9-loxP-LEU2 loxP-CDC20-Intron-loxP-HPHMX Tom70-GFP-TRP1/+ RPL13A-mCherry-Kan/+* |
| --- | --- | --- |
| JH1551 | **ACC1^S1157A^-RPL13a**  MEP Tom70-GFP Rpl13a-mCherry acc1-S1157A het. | ade2::hisG his3 leu2 met15D::ADE2/+ lys2/+ ura3D0 trp1D63 hoD::SCW11pr-Cre-EBD78-NatMX loxP-UBC9-loxP-LEU2 loxP-CDC20-Intron-loxP-HPHMX Tom70-GFP-TRP1/+ RPL13A-mCherry-Kan/+ acc1-S1157A/+ |
| DH83 | **P_GPD_-SAK1-RPL13a**  MEP diploid TOM70-GFP RPL13A-mCherry SAK1 oe | ade2::hisG his3 leu2 met15D::ADE2 ura3D0 trp1D63 hoD::SCW11pr-Cre-EBD78-NatMX loxP-UBC9-loxP-LEU2 loxP-CDC20-Intron-loxP-HPHMX TOM70-GFP-TRP1 RPL13A-mCherry-KanMX6 KanMX6-Pgdp-SAK1 |
| JH1552 | **A2A-RPL13a**  MEP Tom70-GFP Rpl13a-mCherry SAK1 oe acc1-S1157A het. | ade2::hisG his3 leu2 met15D::ADE2/+ lys2/+ ura3D0 trp1D63 hoD::SCW11pr-Cre-EBD78-NatMX loxP-UBC9-loxP-LEU2 loxP-CDC20-Intron-loxP-HPHMX Tom70-GFP-TRP1/+ RPL13A-mCherry-Kan/+ KanMX6-Pgdp-SAK1/+ acc1-S1157A/+ |
| DH192 | **COX9Δ**  MEP diploid TOM70-GFP RPL13A-mCherry cox9Δ | ade2::hisG his3 leu2 met15D::ADE2 ura3D0 trp1D63 hoD::SCW11pr-Cre-EBD78-NatMX loxP-UBC9-loxP-LEU2 loxP-CDC20-Intron-loxP-HPHMX TOM70-GFP-TRP1 RPL13A-mCherry-KanMX6 cox9::URA3 |
| MU19 | **COX9Δ P_GPD_-SAK1**  MEP diploid COX9Δ SAK1oe (TOM70-GFP RPL13A-mCherry) | ade2::hisG his3 leu2 met15D::ADE2 ura3D0 trp1D63 hoD::SCW11pr-Cre-EBD78-NatMX loxP-UBC9-loxP-LEU2 loxP-CDC20-Intron-loxP-HPHMX Tom70-GFP-TRP1 RPL13A-mCherry-Kan cox9::URA3 KanMX6-Pgdp-SAK1 |
| JH1676 | **COX9Δ A2A**  cox9Δ/cox9Δ SAK1 oe acc1-S1157A Tom70-GFP Rpl13-mCherry het dip | ade2::hisG his3 leu2 lys2/+ met15::ADE2/+ ura3DO trp1D63 hoD::SCW11pr-Cre-EBD78-NatMX loxP-UBC9-loxP-LEU2 loxP-CDC20-Intron-loxP-HPHMX KanMX6-Pgdp-SAK1/+ acc1-S1157A/+ cox9::URA3/cox9::URA3 TOM70-GFP-Kan/+ RPL13a-mCherrry-TRP1/+ |
| MU38 | **MPC1Δ**  MPC1Δ MEP diploid (TOM70-GFP VPH1-mCherry) | ade2::hisG his3 leu2 met15D::ADE2 ura3D0 trp1D63 hoD::SCW11pr-Cre-EBD78-NatMX loxP-UBC9-loxP-LEU2 loxP-CDC20-Intron-loxP-HPHMX Tom70-GFP-TRP1 VPH1-mCherry-Kan MPC1::HIS |
| MU39 | **MPC1Δ P_GPD_-SAK1**  MPC1Δ SAK1 oe MEP diploid (TOM70-GFP VPH1-mCherry) | ade2::hisG his3 leu2 met15D::ADE2 ura3D0 trp1D63 hoD::SCW11pr-Cre-EBD78-NatMX loxP-UBC9-loxP-LEU2 loxP-CDC20-Intron-loxP-HPHMX Tom70-GFP-TRP1 VPH1-mCherry-Kan KanMX6-Pgdp-SAK1 MPC1::HIS |
| MU40 | **MPC1Δ A2A**  MPC1Δ + A2A MEP diploid (TOM70-GFP VPH1-mCherry) | ade2::hisG his3 leu2 met15D::ADE2 ura3D0 trp1D63 hoD::SCW11pr-Cre-EBD78-NatMX loxP-UBC9-loxP-LEU2 loxP-CDC20-Intron-loxP-HPHMX Tom70-GFP-TRP1 VPH1-mCherry-Kan KanMX6-Pgdp-SAK1 acc1-S1157A MPC1::HIS |
| HH40HH40 | **MLS1Δ**  MLS1 Δ MEP diploid (Tom70-GFP RPL13A-mCherry) | MAT α ade2::hisG his3 leu2 met15D::ADE2 ura3D0 trp1D63 hoD::SCW11pr-Cre-EBD78-NatMX loxP-UBC9-loxP-LEU2 loxP-CDC20-Intron-loxP-HPHMX Tom70-GFP-TRP1 RPL13A-mCherry-Kan MLS1::URA3 |
| MU31 | **MLS1Δ P_GPD_-SAK1**  MLS1Δ SAK1 oe MEP diploid (Tom70-GFP RPL13A-mCherry) | ade2::hisG his3 leu2 lys2 ura3DO trp1D63 hoD::SCW11pr-Cre-EBD78-NatMX loxP-UBC9-loxP-LEU2 loxP-CDC20-Intron-loxP-HPHMX KanMX6-Pgdp-SAK1 MLS1::URA3 Tom70-GFP-TRP1 RPL13A-mCherry-Kan |
| MU59 | **MLS1Δ A2A**  MLS1 Δ + Sak1 oe + ACC1 S1157A MEP diploid (Tom70-GFP RPL13A-mCherry) | ade2::hisG his3 leu2 met15D::ADE2 ura3D0 trp1D63 hoD::SCW11pr-Cre-EBD78-NatMX loxP-UBC9-loxP-LEU2 loxP-CDC20-Intron-loxP-HPHMX Tom70-GFP-TRP1 RPL13A-mCherry-Kan KanMX6-Pgdp-SAK1 acc1-S1157A MLS1::URA3 |
| HH47 | **SFC1Δ**  SFC1Δ MEP diploid (TOM70-GFP + Rpl13a-mCherry) | MAT α ade2::hisG his3 leu2 met15D::ADE2 ura3D0 trp1D63 hoD::SCW11pr-Cre-EBD78-NatMX loxP-UBC9-loxP-LEU2 loxP-CDC20-Intron-loxP-HPHMX Tom70-GFP-TRP1 RPL13A-mCherry-Kan SFC1::URA3 |
| MU32 | **SFC1Δ P_GPD_-SAK1**  SFC1Δ SAK1 oe MEP diploid (Tom70-GFP RPL13A-mCherry) | ade2::hisG his3 leu2 lys2 ura3DO trp1D63 hoD::SCW11pr-Cre-EBD78-NatMX loxP-UBC9-loxP-LEU2 loxP-CDC20-Intron-loxP-HPHMX KanMX6-Pgdp-SAK1 SFC1::URA3 Tom70-GFP-TRP1 RPL13A-mCherry-Kan |
| MU66 | **CAT2Δ MLS1Δ**  CAT2Δ + MLS1 Δ MEP diploid (Tom70-GFP RPL13A-mCherry) | ade2::hisG his3 leu2 lys2 ura3DO trp1D63 hoD::SCW11pr-Cre-EBD78-NatMX loxP-UBC9-loxP-LEU2 loxP-CDC20-Intron-loxP-HPHMX Tom70-GFP-TRP1 RPL13A-mCherry-Kan MLS1::URA3 CAT2::HIS |
| MU45 | **CAT2Δ MLS1Δ P_GPD_-SAK1**  CAT2Δ + MLS1Δ + SAK1o/e MEP diploid (Tom70-GFP RPL13A-mCherry) | ade2::hisG his3 leu2 met15D::ADE2 ura3D0 trp1D63 hoD::SCW11pr-Cre-EBD78-NatMX loxP-UBC9-loxP-LEU2 loxP-CDC20-Intron-loxP-HPHMX Tom70-GFP-TRP1 RPL13A-mCherry-Kan KanMX6-Pgdp-SAK1 MLS1::URA3 CAT2::HIS |
| MU69 | **CAT2Δ MLS1Δ A2A**  CAT2Δ MLS1Δ SAK1 oe acc1-S1157A MEP diploid (Tom70-GFP RPL13A-mCherry) | ade2::hisG his3 leu2 lys2 ura3DO trp1D63 hoD::SCW11pr-Cre-EBD78-NatMX loxP-UBC9-loxP-LEU2 loxP-CDC20-Intron-loxP-HPHMX KanMX6-Pgdp-SAK1 acc1-S1157A MLS1::URA3 CAT2::HIS Tom70-GFP-TRP1 RPL13A-mCherry-Kan |
| HH88 | **SIP2Δ**  Sip2Δ MEP diploid TOM70-GFP VPH1-mCherry | MAT α ade2::hisG his3 leu2 met15D::ADE2 ura3D0 trp1D63 hoD::SCW11pr-Cre-EBD78-NatMX loxP-UBC9-loxP-LEU2 loxP-CDC20-Intron-loxP-HPHMX Tom70-GFP-TRP1 VPH1-mCherry-Kan Sip2::URA3 |
| HH108 | **SIP2Δ P_GPD_-SAK1**  Sip2Δ MEP diploid SAK1 oe (Tom70-GFP + VPH1-mCherry) | MAT a ade2::hisG his3 leu2 lys2 ura3DO trp1D63 hoD::SCW11pr-Cre-EBD78-NatMX loxP-UBC9-loxP-LEU2 loxP-CDC20-Intron-loxP-HPHMX KanMX6-Pgdp-SAK1 Sip2::URA3 Tom70-GFP-TRP1 VPH1-mCherry-Kan |
| HH87 | **SIP2Δ A2A**  Sip2Δ SAK1 oe acc1-S1157A (Tom70-GFP + VPH1-mCherry) | MAT a ade2::hisG his3 leu2 lys2 ura3DO trp1D63 hoD::SCW11pr-Cre-EBD78-NatMX loxP-UBC9-loxP-LEU2 loxP-CDC20-Intron-loxP-HPHMX KanMX6-Pgdp-SAK1 acc1-S1157A Sip2::URA3 Tom70-GFP-TRP1 VPH1-mCherry-Kan |
| MU64 | **CAT2Δ**  MEP diploid (Tom70-GFP RPL13A-mCherry) | ade2::hisG his3 leu2 met15D::ADE2 ura3D0 trp1D63 hoD::SCW11pr-Cre-EBD78-NatMX loxP-UBC9-loxP-LEU2 loxP-CDC20-Intron-loxP-HPHMX Tom70-GFP-TRP1 RPL13A-mCherry-Kan CAT2::HIS |
